# Systematic discovery and functional interrogation of SARS-CoV-2 viral RNA-host protein interactions during infection

**DOI:** 10.1101/2020.10.06.327445

**Authors:** Ryan A. Flynn, Julia A. Belk, Yanyan Qi, Yuki Yasumoto, Cameron O. Schmitz, Maxwell R. Mumbach, Aditi Limaye, Jin Wei, Mia Madel Alfajaro, Kevin R. Parker, Howard Y. Chang, Tamas L. Horvath, Jan E. Carette, Carolyn Bertozzi, Craig B. Wilen, Ansuman T. Satpathy

## Abstract

Severe acute respiratory syndrome coronavirus 2 (SARS-CoV-2) is the cause of a pandemic with growing global mortality. There is an urgent need to understand the molecular pathways required for host infection and anti-viral immunity. Using comprehensive identification of RNA-binding proteins by mass spectrometry (ChIRP-MS), we identified 309 host proteins that bind the SARS-CoV-2 RNA during active infection. Integration of this data with viral ChIRP-MS data from three other positive-sense RNA viruses defined pan-viral and SARS-CoV-2-specific host interactions. Functional interrogation of these factors with a genome-wide CRISPR screen revealed that the vast majority of viral RNA-binding proteins protect the host from virus-induced cell death, and we identified known and novel anti-viral proteins that regulate SARS-CoV-2 pathogenicity. Finally, our RNA-centric approach demonstrated a physical connection between SARS-CoV-2 RNA and host mitochondria, which we validated with functional and electron microscopy data, providing new insights into a more general virus-specific protein logic for mitochondrial interactions. Altogether, these data provide a comprehensive catalogue of SARS-CoV-2 RNA-host protein interactions, which may inform future studies to understand the mechanisms of viral pathogenesis, as well as nominate host pathways that could be targeted for therapeutic benefit.

**Highlights:**

· ChIRP-MS of SARS-CoV-2 RNA identifies a comprehensive viral RNA-host protein interaction network during infection across two species
· Comparison to RNA-protein interaction networks with Zika virus, dengue virus, and rhinovirus identify SARS-CoV-2-specific and pan-viral RNA protein complexes and highlights distinct intracellular trafficking pathways
· Intersection of ChIRP-MS and genome-wide CRISPR screens identify novel SARS-CoV-2-binding proteins with pro- and anti-viral function
· Viral RNA-RNA and RNA-protein interactions reveal specific SARS-CoV-2-mediated mitochondrial dysfunction during infection

## Introduction

Despite similarities in replication strategies of their compact genomes, positive single stranded RNA viruses cause a remarkable variety of human diseases. Mosquito-borne flaviviruses such as dengue virus and Zika virus cause systemic disease, while human coronaviruses generally cause respiratory symptoms (Ahlquist, 2006; Carrasco-Hernandez et al., 2017). The recent pandemic emergence of the novel coronavirus SARS-CoV-2, which can cause potentially fatal COVID-19 disease, illustrates the threat to public health posed by RNA viruses. Less than one year into the outbreak, more than 33 million people have been infected by SARS-CoV-2, and one million people have died. The severity of the virus has caused global economic disruption and treatment options remain limited, due to, in part, an incomplete understanding of the molecular determinants of viral pathogenesis.

The process of infecting a host cell is complex, multistep, and often highly virus-specific. Viruses must traffic to, bind, and enter host cells, and once inside the cell, their genetic material leverages and remodels cellular pathways to express, replicate, and produce new infectious virions. RNA viruses deposit large autonomous RNA transcripts into the dense intracellular milieu of the host cells, which eventually generate virally-encoded protein products. Together, these RNA and protein species remodel the cell to facilitate the viral life cycle. We and others have demonstrated the utility of functionally exploring how different virally-derived molecules hijack the host, in particular in the context of flaviviruses (Li et al., 2020). For example, mapping physical associations between the host and virus at the level of protein-protein interactions (PPI) have defined key pathways relevant to infection (Eckhardt et al., 2020). In parallel to efforts that focus on viral proteins, a number of groups have taken an RNA-centric view of the host-viral interface to understand how host cells recognize and respond to the RNA genome (Kim et al., 2020a; Lenarcic et al., 2013; Ooi et al., 2019; Phillips et al., 2016; Viktorovskaya et al., 2016). Finally, genetic screening efforts provide another, more direct strategy to discover cellular proteins and pathways that are essential for viral replication or that are part of the host innate immune responses (Puschnik et al., 2017; Schoggins and Rice, 2011). Each of these approaches has limitations and captures only one aspect of the host-virus interface, but together they begin to unravel the complex machinery evolved by each virus to invade and function within the host cell.

While there has been significant past work to understand coronaviruses (Cockrell et al., 2018; Gralinski and Baric, 2015), the emergence of novel strains which are highly transmissible and cause severe disease in humans (Menachery et al., 2015) has underscored the need to refine our understanding of: (1) the basic mechanisms of their life cycle, (2) how they modulate host processes at both the cellular and organismal level, and (3) how the host combats infection with intracellular and systems-level defense mechanisms. Recent studies have described SARS-CoV-2 encoded proteins (Kim et al., 2020b) and their interacting host partners (Gordon et al., 2020); however, there is a gap in understanding of the precise host interactions of the SARS-CoV-2 viral RNA (vRNA). To address this gap, we used ChIRP-MS (Chu et al., 2015), which provides a comprehensive view of the host interactions of vRNAs spanning all subcellular domains where the RNA is present. This strategy provided an opportunity to define the shared and SARS-CoV-2-specific host pathways that associate with vRNAs. We combined the RNA-centric approach with genome-wide genetic perturbations to define known and novel host factors relevant to the SARS-CoV-2 life cycle and discovered a functional and specific interface between SARS-CoV-2 and the host mitochondria.

## Results

### ChIRP-MS of SARS-CoV-2 viral RNA in infected mammalian cells

To define the host protein interactome of the ∼30kb SARS-CoV-2 vRNA, we performed ChIRP-MS (**Figure 1A**). ChIRP-MS is advantageous as a discovery tool because it uses formaldehyde as a crosslinking agent to recover entire protein complexes associated with cellular RNAs (Chu and Chang, 2018; Chu et al., 2015). We selected two cell lines: (1) Huh7.5, one of the few human cell models that is naturally susceptible to productive infection by SARS-CoV-2, and (2) VeroE6, a monkey kidney cell line that dominates the research space for preparation and propagation of SARS-CoV-2 and other viruses (Harcourt et al., 2020; Zhou et al., 2020). We tiled 108 biotinylated oligonucleotide probes (**Table S1**) to capture the full length positive-strand vRNA, which includes subgenomic RNA (sgRNA) species that accumulate to higher copy numbers during infection (Kim et al., 2020b). In addition to the interspecies and cell type comparisons, we also performed ChIRP-MS experiments at two different time points, 24- and 48-hours post infection (h.p.i.), to assess the temporal association of host factors with the vRNA (**Figure 1A**). From each condition, input and ChIRP-enriched RNA and protein samples were saved for later analysis (**Figure 1A**). Initially, we assessed the recovery and profile of the enriched proteins by SDS-PAGE analysis (**Figure 1B**). As expected, mock samples (uninfected, acting as probe-only controls) had little protein staining, while we observed an infection and time-dependent increase in total protein recovered after infection of either cell line with SARS-CoV-2 (**Figure 1B**). The clear band present in all infected samples at ∼50kDa is consistent with the viral nucleocapsid (N) protein (**Figure 1B**), a major RNA binding protein (RBP) of the coronavirus RNAs (Chang et al., 2014). We next assessed the technical quality of the ChIRP experiment by analyzing viral and host RNAs recovered before and after pulldown. RNA sequencing from mock samples resulted in negligible mapping to the SARS-CoV-2 genome before or after pulldown, as expected (**Figure S1A**). In contrast, in SARS-CoV-2 infected cells, we observed a basal level of 2.7% (Huh7.5 48 h.p.i) and 14.4% (VeroE6 48 h.p.i) of all reads in total RNA mapping to the viral genomic RNA, which increased to 60% (Huh7.57 48 h.p.i) and 68% (VeroE6 48 h.p.i) after pulldown, demonstrating robust enrichment of vRNA after ChIRP (**Figure S1A**). These results are consistent with prior studies demonstrating higher rates of infection in VeroE6 cells, compared to most other cell lines (Harcourt et al., 2020). Since coronaviruses produce full length, as well as sgRNAs, we next assessed whether the ChIRP-MS was biased for the higher molar copy sgRNAs. Analysis of the reads mapping to the SARS-CoV-2 genomic RNA showed that both input and ChIRP-enriched samples had robust coverage of the ORF1a/b region, as well as in the sgRNA regions, and that the enrichment was visually and quantitatively similar across Huh7.5 and VeroE6 (**Figure 1C-E**). Together these protein- and RNA-level quality controls demonstrate the robust and broad sampling of the entire SARS-CoV-2 positive-strand vRNA by the designed ChIRP-MS probes.

**Figure 1:**
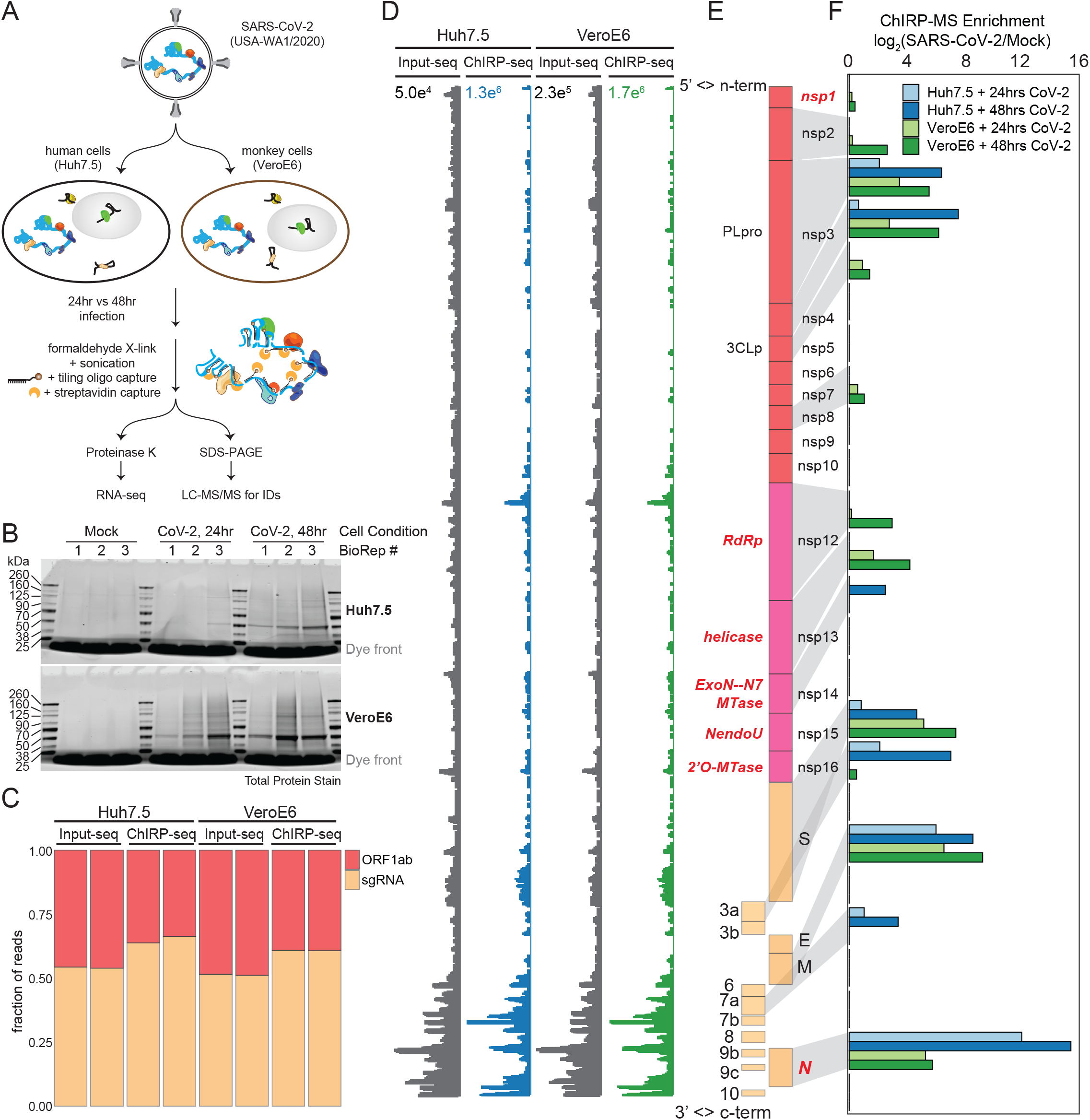
ChIRP-MS identifies host and viral proteins associated with the SARS-CoV-2 RNA genome in infected cells. **(A)** Schematic of the ChIRP-MS protocol. **(B)** SDS-PAGE analysis of total protein samples enriched using SARS-CoV-2 targeting biotinylated oligonucleotides from Mock (uninfected) cells or cells infected for 24 or 48 hours with SARS-CoV-2 in both Huh7.5 (top) and VeroE6 (bottom) cells. **(C)** Quantification of the percentage of reads mapping to SARS-CoV-2 gRNA (ORF1a/b) versus the subgenomic RNA (sgRNA) before and after pulldown. **(D)** RNA-seq coverage of the SARS-CoV-2 genome before and after pulldown. **(E)** Structure of the SARS-CoV-2 genome. **(F)** Enrichment of each viral protein in Huh7.5 and VeroE6 cells at both time points.

We first asked whether any of the SARS-CoV-2 encoded proteins were enriched after pulldown of the vRNA. Viruses hijack host factors but also encode a select group of proteins that are critical for their life cycle. To date, SARS-CoV-2 has been shown to encode 16 nonstructural proteins, 4 structural proteins and 6 accessory proteins (Finkel et al., 2020), several with annotated RNA binding capacity (**Figure 1E**, known RBPs in red). We observed that 13 of 26 viral proteins were reproducibly enriched (mean log_2_ enrichment ≥ 1 in at least one dataset), including known and novel RNA-binding viral proteins. In the sgRNA region, the major viral proteins conserved across cell types were N, M, and S, while ORF3a and 7a were selectively enriched from infected Huh7.5 cells (**Figure 1F**). Within the larger ORF1a/b, nsp3 and nsp4 were enriched in both species, however we saw stronger association of the annotated RBPs in VeroE6 infected cells (**Figure 1F**). The robust enrichment of specific ORF1a/b encoded proteins provides strong evidence that the ChIRP-MS approach samples interactions across the entire length of the genomic RNA. Considering the VeroE6- and Huh7.5-specific associations, we note that these cells have many differences (including species, cell type, and sex of organism), any of which may cause the observed differences in ChIRP enrichments. One consideration is that VeroE6 cells are well characterized to support high SARS-CoV-2 replication and efficient viral egress, while replication in Huh7.5 cells reaches lower peak levels with delayed kinetics (Harcourt et al., 2020). Nonetheless, viral protein enrichments were quite specific and reproducible, and the different features of these cell lines enable us to define a core SARS-CoV-2-associated proteome which is conserved across these cell types and infection timepoints.

### A comprehensive atlas of host-factors that interact with the SARS-CoV-2 genomic RNA

To define the host-derived interacting proteins of the SARS-CoV-2 vRNA, we searched the ChIRP-MS data against a database of known monkey (VeroE6) or human (Huh7.5) proteins. Using stringent statistical cutoffs (*p* adjusted ≤ 0.05, fold change > 0), we defined high-confidence interactomes in each condition (**Figure 2A, 2B**). A total of 163 (VeroE6) and 229 (Huh7.5) host factors bound to the SARS-CoV-2 vRNA (**Table S2**). Analysis of the factors enriched at 24 vs. 48 h.p.i revealed that most factors enriched in VeroE6 cells were invariant between the two time points (**Figure 2C**, left) while the Huh7.5 interactome evolved more dramatically over this period, with 48 h.p.i. showing an expanded set of interacting proteins (**Figure 2C**, middle). We next compared the associated host factors across species and found a core set of 83 factors co-bound in both species, totaling 309 host factors aggregated across species (**Figure 2C**, right). The Huh7.5 cells accumulated the greatest number of specific host factors during the course of infection, and we found the human proteome to be substantially better annotated than the monkey proteome, so we focused our analysis on the human dataset.

**Figure 2:**
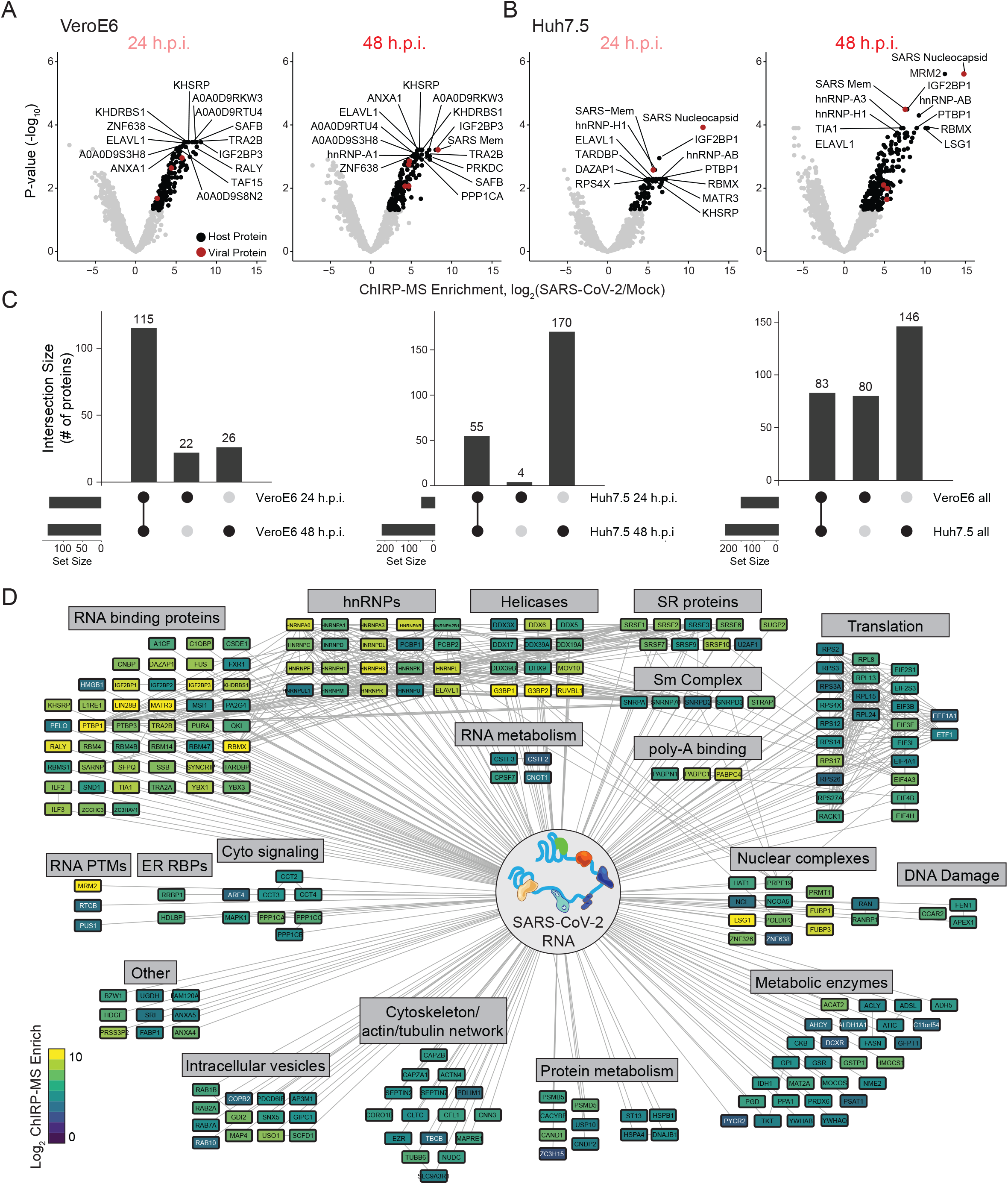
Changes in the SARS-CoV-2 associated proteome across time points and species. **(A)** Significantly enriched proteins in Vero cells after viral RNA pulldown at 24 and 48 h.p.i. **(B)** Significantly enriched proteins in Huh7.5 cells after viral RNA pulldown at 24 and 48 h.p.i. **(C)** Conservation of enriched proteins between time points (left, middle) and species (right). **(D)** Cytoscape network representation of the SARS-CoV-2 associated human proteome. Colors indicate ChIRP enrichment in Huh7.5 cells 48 h.p.i.

In an effort to better understand the set of host factors associated with the SARS-CoV-2 RNA, we visualized the high confidence human interactome using Cytoscape (Shannon et al., 2003) (**Figure 2D**). In this analysis, each node is a protein significantly enriched in the Huh7.5 ChIRP-MS dataset in at least one time point and is colored by its enrichment at 48 h.p.i. in Huh7.5 cells. Nodes are connected if there is a previously described protein-protein interaction (**Figure 2D**). Structuring this network by broad functional categories demonstrated the diversity of host proteins associated with the vRNA, spanning generic RNA adaptor proteins, RNA helicases, RNA processing enzymes, and RNA modification enzymes (**Figure 2D**). We also noticed a set of relatively unexpected pathways which had multiple robust factors binding, including a suite of metabolic enzymes, intracellular vesicle proteins, cytosolic signaling, and cytoskeleton and intracellular trafficking proteins (**Figure 2D**).

Using this network, we next performed a series of comparisons in order to contextualize the ChIRP-MS results. First, we asked how the host viral-binding factors changed from 24 to 48 h.p.i. We found that earlier in the infection in Huh7.5 cells, a core set of RBPs were strongly bound (top left region of the network) while the rest of the network only became associated after 48 hours, suggesting these RBPs may be important for the earliest steps of detection or replication of the vRNA (**Figure S2A**). Comparing the VeroE6 and Huh7.5 interactomes showed that the core RBPs were also very highly conserved across species, as were other categories such as nuclear complexes, poly-A binding proteins, and serine/arginine rich splicing factors (SR proteins) (**Figure S2B**). We next compared the ChIRP-MS results to a set of host factors identified by vRNA pulldown after UV-C crosslinking (RNA Antisense Purification (RAP) MS, Schmidt et al., 2020). We overlaid these proteins onto the ChIRP-MS interaction network and found the majority (30/48, 63%) were also enriched in ChIRP-MS (**Figure S3A, S3B**). However, ChIRP-MS enriched an additional 199 proteins that were not identified as significant in the UV-C dataset. The substantial increase in scope of ChIRP-enriched factors is consistent with prior reports (Ooi et al., 2019) and the broader crosslinking capability of formaldehyde, compared to UV-C, which specifically captures direct RNA-protein interactions. Finally, we compared the ChIRP-MS data to a recently published and comprehensive characterization of the host-viral protein-protein interactome of 26 SARS-CoV-2 encoded proteins (Gordon et al., 2020). We found that only 11/332 host factors (3.3%) from the PPI study overlapped with the ChIRP-MS network (**Figure S3A, S3C**), demonstrating that SARS-CoV-2 vRNA and proteins largely interact with distinct protein complexes inside of the cell. However, of the 11 host factors that bind both vRNA and viral proteins, RAB2A, RAB7A, and RAB10 have been validated as functional in SARS-CoV-2 infection (Hoffmann et al., 2020). Altogether, these comparisons highlight the orthogonality of an RNA-centric approach to PPI-based studies, and the power of formaldehyde crosslinking to discover larger cellular complexes associated with vRNAs during infection.

### Inter-virus analysis of host factors reveals specificity of interacting cellular pathways

It has become increasingly clear that interactions of vRNAs with proteins play key roles in multiple aspects of viral infection, either through the recruitment of host factors essential for viral translation and replication, or as interaction partners for cellular proteins involved in anti-viral responses (Fritzlar et al., 2019; Garcia-Blanco et al., 2016; Hosmillo et al., 2019). To understand commonalities and differences in how positive-stranded RNA viruses have evolved to interact with their host, we sought to compare the SARS-CoV-2 dataset to our previously generated ChIRP-MS data from the flaviviruses Zika (ZIKV, ZIKV-PRVABC59) and Dengue-2 (DENV, DENV-16681), as well as a human picornavirus, rhinovirus (RV, RV-B14) (Ooi et al., 2019). We note that all datasets were collected from Huh7.5 cells except for the rhinovirus data, which was collected from HeLa cells. We first used principal component analysis (PCA) to broadly investigate the associated host factors enriched across viruses and found that PC1 separated all 4 viral types and PC2 further distinguished RV and demonstrated the time-dependent host factor changes for SARS-CoV-2 (**Figure 3A**). Next, to facilitate quantitative comparisons between the viruses, for each virus, we defined an “expanded interactome” consisting of proteins reproducibly enriched (average log_2_ fold change ≥ 1) for each ChIRP-MS dataset in human cells: SARS-D1 (Huh7.5 24 h.p.i.), SARS-D2 (Huh7.5 48 h.p.i.), ZIKV-D2, DENV-D2, and RV-D2. Comparing the datasets at the day 2 time point, each expanded interactome consisted of about 1000 proteins associated with each vRNA (**Figure 3B**). We found that the largest group of 425 proteins was shared across all ChIRP-MS datasets: SARS-D2, both flaviviruses, and RV-D2 (**Figure 3B**). Despite the difference in input proteome, the RV-D2 expanded interactome had a similar number of unique proteins (181) as that of SARS-D2 (167). There were 138 proteins interacting with both flaviviruses (DENV and ZIKV) but neither SARS-CoV-2 nor RV. Full expanded interactome data is provided in **Table S3**.

**Figure 3:**
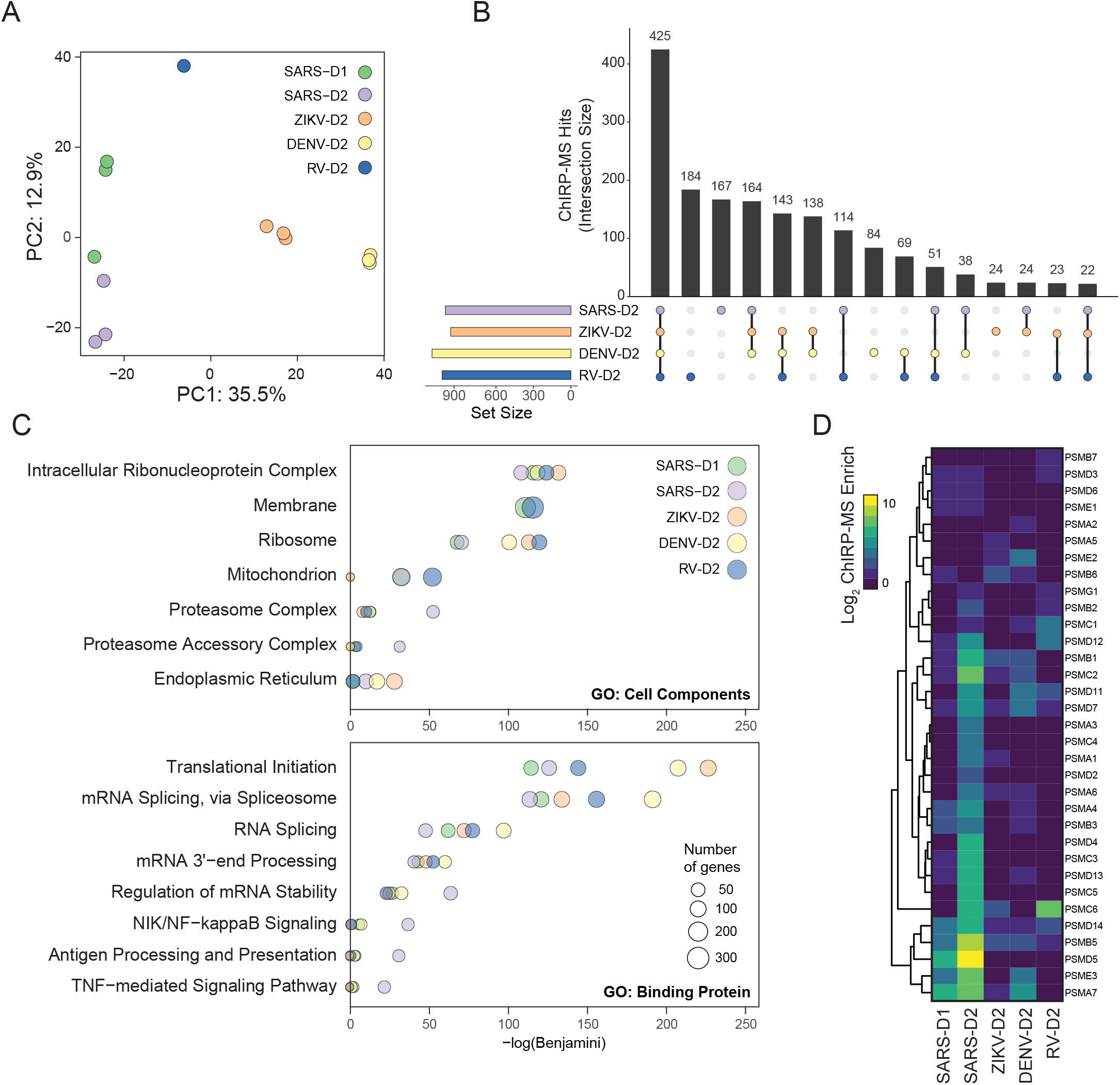
Comparison of the SARS-CoV-2 associated proteome to that of other RNA viruses. **(A)** Principal component analysis of ChIRP enrichments in human cells across time points and viruses. **(B)** Upset plot comparing expanded interactomes of SARS-CoV-2, ZIKV, DENV, and RV in human cells at 48 h.p.i. **(C)** Top: Cellular Components GO terms enriched in the expanded interactome of each virus. Bottom: Binding Protein GO terms enriched in the expanded interactome of each virus. **(D)** Comparison of proteasome subunits and proteasome accessory factor associations across viruses.

To understand the general pathways and associations of each virus, we performed GO term analysis on the expanded interactomes of each virus. Consistent with the RNA-centric aspect of ChIRP, all viruses robustly enriched the intracellular RNP complex term; however, we found patterns of specificity when examining other cellular pathway terms (**Figure 3C**). For example, SARS-CoV-2 displayed a reduced enrichment with the ER and ribosome, but an increased enrichment with mitochondria and proteasome (**Figure 3C**, top). Examining functional terms again corroborated a decreased enrichment with translation and splicing factors in the SARS-CoV-2 interactome, compared to that of the flaviviruses, but a specific increased enrichment with multiple immune pathways such as antigen presentation, NF-κb signaling, and TNF signaling (**Figure 3C**, bottom). While the GO terms provide an initial high level assessment of the associated factors, we next wanted to understand how these pathways or protein complexes look at the individual factor level. Guided by the SARS-CoV-2-specific enrichment of the proteasomal GO term, we visualized all the individual subunits of the proteasome found in any of the ChIRP-MS datasets (**Figure 3D**). Consistent with the GO term analysis, DENV, ZIKV, and RV all poorly associated with this set of proteins while there was broad and robust enrichment with the SARS-CoV-2 vRNA, in particular at D2 (**Figure 3D**). Previous work has reported a functional connection to proper proteasome function and coronavirus life cycles (Raaben et al., 2010), which together with our ChIRP-MS data may suggest the vRNA directly leverages the proteasome during infection, potentially to modulate antigen presentation and/or evade host adaptive immunity. The specificity of association between the proteasome and the SARS-CoV-2 RNA and clear validation of this interaction in the literature motivated us to explore the set of RNA-centric viral interactomes across a number of other important cellular pathways (**Table S3**).

#### RNA binding proteins

First we focused on the heterogeneous nuclear ribonucleoproteins (hnRNPs), a large set of adaptor proteins, and dead-box helicases, which remodel RNA structural elements (Geuens et al., 2016; Jankowsky, 2011). These RBP families have a wide array of cellular functions and are often recovered with host or pathogenic RNAs in binding studies (Geuens et al., 2016; Meier-Stephenson et al., 2018; Taschuk and Cherry, 2020). The hnRNP class showed robust interaction with all 4 viruses and similar enrichments for the majority of the 20 proteins we identified (**Figure 4A**). The DDX proteins, despite being ubiquitously and highly expressed, showed a more virus-specific binding profile wherein family members such as DDX3X, 5, 6, and 38B were similar across viruses, while DDX21, 23, 42, and 46 were more specifically associated with the DENV and RV RNAs (**Figure 4A**). As noted above, these direct RBPs associate with the virus early in infection and may include the initial sensors of infection.

**Figure 4:**
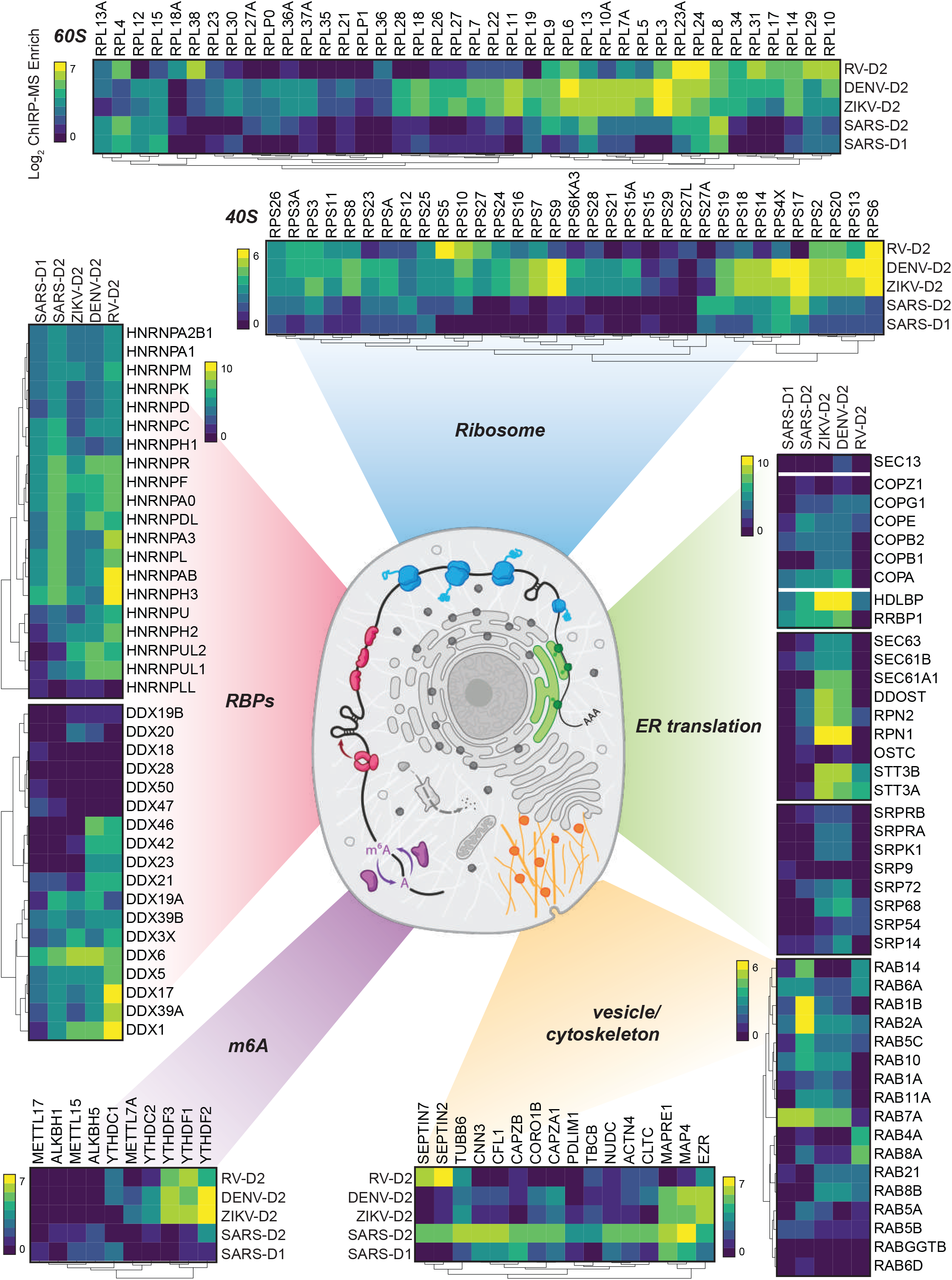
Cellular context of expanded interactomes across viruses. Selected groups of proteins, their enrichment in SARS-CoV-2, Zika, Dengue, and Rhinovirus ChIRP, and their approximate subcellular localization. Heat map colors indicate the log_2_ ChIRP-MS enrichment values. Each heatmap has a separate scale bar.

#### Translational apparatus

After entry into the cytosol, one of the first steps of the viral life cycle is to begin to express the protein products encoded in its genome, which requires interactions with the host translational apparatus. Work examining the translational capacity of RNA viruses has show that, in contrast to flaviviruses, which are more efficiently translated than cellular mRNAs under cellular stress (Edgil et al., 2006; Roth et al., 2017), the coronaviruses do not translate their mRNAs at higher efficiency than cellular mRNAs during infection (Finkel et al., 2020). A comparison of enriched translation initiation factors (eIFs) demonstrated quantitative differences across the viruses: flaviviruses strongly enriched EIF3A, 4G1, 3C and 3D while SARS-CoV-2 was relatively depleted for these factors but preferred EIF3B, 4H, 4B, 3F, A3, among others (**Figure 3C**). These data point to a more specialized configuration of the initiation complex on different viral genomes. Beyond translational initiation, we visualized enrichment for the core components of the 80S ribosome (**Figure 4B**). Here we note that while there was specificity in the enrichment of specific ribosomal proteins (RPs), more striking was the generalized lack of association of the vast majority of the RPs to the SARS-CoV-2 vRNA, compared to either DENV or ZIKV (**Figure 4B**). This is consistent with recent reports demonstrating global translation inhibition by SARS-CoV-2 encoded nsp1 (Schubert et al., 2020; Thoms et al., 2020). How SARS-CoV-2 simultaneously inhibits host translation to shut down innate immune responses yet also manages to translate its own proteins remains unclear, although recent work points to a functional role for the 5’ UTR of the viral RNA--likely together with yet-unidentified translation initiation factors--in promoting efficient translation even in the presence of nsp1 (Schubert et al., 2020).

#### Sec, translocon, and ER-golgi transport

Flaviviruses and coronaviruses encode glycoproteins, but picornaviruses do not. We and others previously showed that RV weakly enriches factors related to membrane biology, in contrast to the functional use of membrane organelles like the ER by flaviviruses (Fernandez-Garcia et al., 2009; Mukhopadhyay et al., 2005; Ooi et al., 2019). Given the strong dependence of flaviviruses on the translocon, the channel for nascent peptide entry into the ER lumen, and adjacent protein complexes, we compared these factors across the viral ChIRP-MS data (**Figure 4C**). We found that while SARS-CoV-2 does robustly enrich ER-tethered (RRBP1) or associated (HDLBP/vigilin) RBPs, it is less strongly associated with the ER-targeting complex (SRP) or the Sec/translocon itself. However, SARS-CoV-2’s vRNA associates with the COPI vesicle complexes in a more similar manner to the flaviviruses. COPI proteins are canonically responsible for retrograde transport of vesicles from the golgi to ER (Szul and Sztul, 2011). The association with COPI complex members is consistent with the reported cycling of SARS-CoV in the ER-Golgi network for eventual budding into the lumen ER-Golgi intermediate compartment (ERGIC, (McBride et al., 2007)). Our data therefore suggest that ER-resident RBPs may be commonly leveraged for flavivirus and coronavirus life cycles, while other membrane-associated factors are more virus-specific.

#### Intracellular vesicles and trafficking

Given the differences in associations with the major translational and translocation machinery at the ER, we explored how other intracellular vesicle and trafficking complexes might differ across viruses. This is of additional interest in SARS-CoV-2 infection given the developing evidence of the intracellular double-membrane vesicles which are produced (Wolff et al., 2020). We found many host factors involved in cytokinesis, actin filaments, cytoskeleton, and microtubules associated with the vRNAs but a particular bias in association with the SARS-CoV-2 vRNA relative to others (**Figure 4D**). Recent reports highlighted the physical association of Rab GTPase family members with viral proteins and their functional importance in the temperature-dependent life cycle of coronaviruses (Gordon et al., 2020; Hoffmann et al., 2020). Our ChIRP-MS data robustly supports these observations as we find that four Rab proteins, RAB1B, RAB2A, RAB7A, and RAB10, were all present in the SARS-CoV-2 high confidence interactome (**Figure 2D**) with multiple others strongly associated with the SARS-CoV-2 RNA (**Figure 4D**). Looking beyond SARS-CoV-2, we found that DENV and ZIKV also strongly recover RAB2A and RAB10, which could suggest these viruses may be subject to similar regulation in this pathway as SARS-CoV-2.

#### N^6^-methyladenosine

Post-transcriptional regulation of RNA is a rapidly growing field of study and one chemical modification that has received renewed interest has been m6A: methylation of the N-6 position on adenine (Yue et al., 2015; Zaccara et al., 2019). A wide range of cellular processes have been now associated with this modification and recently it has been reported that m6A is deposited on the vRNA of ZIKV (Lichinchi et al., 2016). This appears to be an anti-viral mechanism as binding of YTH-family proteins (which recognize m6A) subsequently cause degradation of the ZIKV vRNA (Lichinchi et al., 2016). We therefore examined the association of the writers (METTL-family), readers (YTH-family) and erasers (ALKBH-family) of m6A across the 4 viral ChIRP-MS datasets. Consistent with the work reporting m6A’s role in ZIKV infection, we saw a robust association of the YTHDF-family with ZIKV and DENV genomic RNAs (**Figure 4E**). RV also robustly captured these proteins however SARS-CoV-2 lacked robust enrichment of these proteins. Conversely, we found relatively stronger enrichment of the m6A-demethylases associated with the SARS-CoV-2 vRNA while ZIKV, DENV, and RV all poorly bound these proteins (**Figure 4E**). ChIRP-MS therefore suggests that SARS-CoV-2 may evade robust m6A modification in order to stabilize it’s vRNA inside infected cells.

### Intersection of ChIRP-MS and CRISPR perturbation screens assign functional relevance to RNA-protein interactions

To understand the functional role of host proteins identified by ChIRP-MS in SARS-CoV-2 pathogenicity, we intersected the ChIRP-MS results with CRISPR perturbation data from our previous study, which utilized a library composed of 83,963 gene-targeting single guide RNAs to identify host genes essential for cell survival after SARS-CoV-2 infection in VeroE6 cells (Wei et al., 2020). This CRISPR screen was designed to identify both pro- and anti-viral host factors: depletion of guide RNAs after infection indicated that the gene had host-protective or anti-viral function, and knockout of the gene sensitized the cell to virus-induced cell death, while enrichment of guide RNAs after infection indicated that the gene had pro-viral function, and knockout of the gene conferred resistance to virus-induced cell death. We first calculated CRISPR z-scores for the core (309) and expanded (1430) host protein interactomes identified by ChIRP-MS and identified 33 core factors and 98 expanded factors that had a functional impact on host cell survival after SARS-CoV-2 infection (**Figure 5A-D**). Surprisingly, CRISPR-targeting of 29/33 core factors and 87/98 expanded factors resulted in sensitization to SARS-CoV-2-induced cell death, demonstrating that the vast majority of host factors that bind the vRNA are host-protective, rather than pro-viral factors. For both interactomes, this bias towards sensitizing genes was significant compared to the distribution of all hits in the genome-wide screen (p < 0.0002, all hits vs core ChIRP-MS interactome, and p < 2×10^−7^, all hits vs expanded ChIRP-MS interactome, Mann-Whitney test, **Figure 5C**).

**Figure 5:**
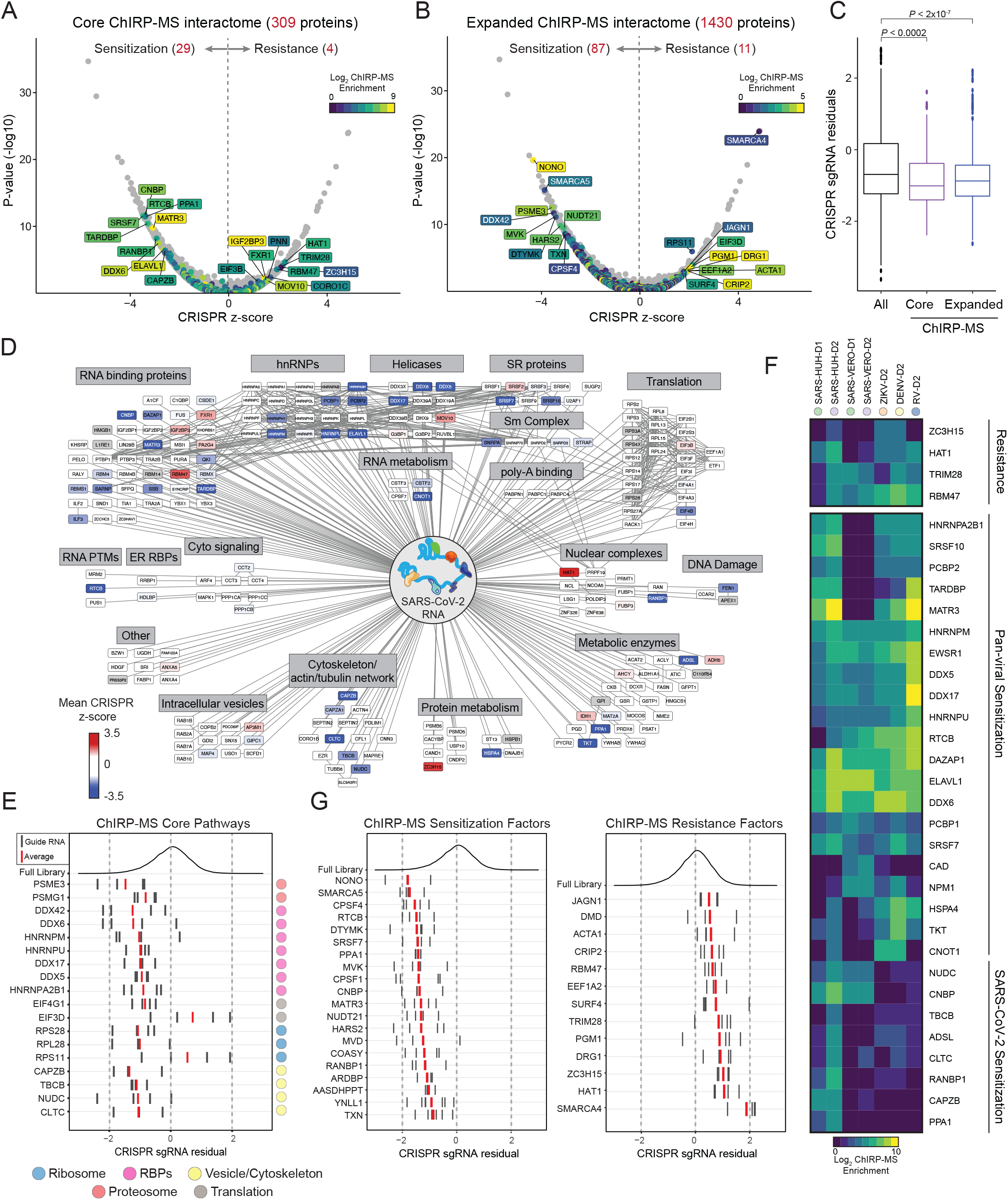
Integration of ChIRP-MS and functional genomic data suggest novel pro- and anti-viral host factors. **(A)** High confidence SARS-CoV-2 interactome overlaid on VeroE6 CRISPR screen data. **(B)** Expanded SARS-CoV-2 interactome overlaid on CRISPR screen data. **(C)** Comparison of CRISPR guide RNA (sgRNA) residuals for significant hits (fdr <= 0.05) of all genes (left, black), genes present in the high-confidence SARS-CoV-2 RNA interactome (purple, middle), or genes present in the expanded SARS-CoV-2 RNA interactome (right, blue). P values computed from Mann-Whitney test. **(D)** High confidence SARS-CoV-2 human interactome network colored by enrichment or depletion in CRISPR screen. **(E)** sgRNA residuals for CRISPR hits identified in (B) grouped by cellular pathways in Figure 4. Individual CRISPR guides are represented by black lines. The average of these is shown in red. **(F)** Inter-virus comparison of shared ChIRP-MS / CRISPR hits identified in (A). **(G)** sgRNA residuals for top 20 sensitizing hits (left) and for all significant resistance hits (right) identified in (B).

Further examination of the functional hits in the SARS-CoV-2 ChIRP-MS interactome revealed known and novel genes in viral pathogenesis (**Figure 5A-D**). For example, TARDBP knockout sensitizes cells to SARS-CoV-2 infection, and indeed, this protein has previously been shown to have anti-viral activity in the context of Human Immunodeficiency Virus type 1 (HIV-1) infection in humans by directly binding to a particular regulatory motif within the HIV-1 RNA genome and thereby repressing viral gene expression (Ou et al., 1995). Similarly, NONO has a previously characterized anti-viral role in the context of HIV and poliovirus infection, where it acts as a sensor for viral DNA and activates cGAS to trigger innate immunity (Lahaye et al., 2018; Lenarcic et al., 2013). CNBP is another factor that sensitizes host cells to infection and has also been shown to recognize diverse intracellular microbial products to drive inflammatory cytokine gene expression, particularly IL-12β (Chen et al., 2018). Members of the SWI/SNF protein family, SMARCA4 and SMARCA5 were among the strongest genome-wide hits in either direction (Wei et al., 2020). SWI/SNF proteins can form multiple distinct complexes and are thought to function predominantly in the nucleus, but we identified associations with the vRNA. Both SMARCA4 and SMARCA5 are ATPases but participate in distinct protein complexes, consistent with their opposing functional activities (**Figure 5B**). We also recovered SMARCC1 in the ChIRP-MS, consistent with its known complex with SMARCA4. Querying known SMARCA5 interacting partners, we additionally identified BAZ1A (also known as ACF1) as present in the VeroE6 expanded interactome, and BAZ1A depletion also sensitized cells to virus-induced death, although less strongly than SMARCA5, suggesting a direct interaction and functional role for the BAZ1A/SMARCA5 complex in viral infection (Oppikofer et al., 2017, **Figure S5A**).

Using this list of functional SARS-CoV-2 RNA-binding host factors, we again performed a series of comparisons to understand their cellular pathways, localization, and virus specificity. First, we compared the CRISPR data with the previously-described Cytoscape human ChIRP-MS interactome (**Figure 5D**). We observed that the majority of functional hits were RBPs, helicases, and hnRNPs, which we defined as core factors that bind the vRNA early during infection (D1), suggesting that the host’s initial response to viral infection is to mount a diverse vRNA recognition program to restrict the viral life cycle. Extending this analysis in the context of the cellular organelles and localization described in Figure 4, we found significant CRISPR hits spanning multiple (but not all) organelles, including proteasome accessory factors PSME3 and PSMG1, several initiation factors and ribosomal proteins, and cytoskeletal proteins (**Figure 5E**). Again, disruption of most of these genes sensitized cells to SARS-CoV-2 induced cell death; however, there were exceptions such as EIF3D and RPS11, which may indicate that the virus co-opts these factors to preferentially translate viral RNAs. Next, we asked whether functional vRNA-binding factors specifically bound SARS-CoV-2 RNA or were pan-viral factors that could also be observed in ChIRP-MS of flavivirus and rhinovirus RNA. Of 29 core sensitization proteins, 21 (75%) were commonly bound by all viruses, suggesting that there is substantial shared host machinery to sense viral RNA (**Figure 5F**). 8 proteins were SARS-CoV-2-specific, including CNBP, PPA1, and CLTC, which may represent novel pathways that protect host cells from coronavirus-induced cell death. Interestingly, many of the SARS-CoV-2-specific sensitization factors were annotated as having cytoskeletal function, such as CAPZB, NUDC, TBCB, and CLTC, potentially indicating a novel aspect of SARS-CoV-2 pathogenesis relative to the other viruses (**Figure 5D, 5F**). As with core factors, 56/87 (64%) of expanded ChIRP-MS sensitization factors were also commonly bound by all viruses. A comprehensive catalog of these factors, and their association across viruses, is provided as a resource (**Figure S5B** and **Table S3**). Finally, we visualized the top 20 sensitizing hits and the complete list (12 total) of resistance hits present in the expanded interactome (**Figure 5G**). We were particularly interested in the resistance hits, since they may have direct relevance as drug targets. Two of the most protective hits, ZC3H15 (also known as DFRP1) and DRG1 are proteins that have been shown to interact with each other as well as the TNF signaling pathway via TRAF-2, and may represent an as-yet unappreciated vulnerability of SARS-CoV-2 and interface between the virus and host immunity (**Figure 5G**, Capalbo et al., 2013; Glingston et al., 2019). In aggregate, we have identified more than 100 host proteins that directly interact with the SARS-CoV-2 vRNA and are functionally implicated in viral pathogenesis.

### A RNA-centric view of SARS-CoV-2 reveal a specific perturbation of mitochondrial during infection

The ChIRP methodology enriches any cellular biopolymer close enough to be crosslinked with formaldehyde, including not only proteins, but also DNA and RNA. Because of this feature, we re-examined the RNA samples initially purified for quality control (**Figure 1**) and aligned the reads which did not map to the viral genome to the host genome (**Figure S1A**). Differential expression analysis revealed 264 and 167 RNAs significantly enriched with the SARS-CoV-2 vRNA in VeroE6 and Huh7.5 cells, respectively, at 48 h.p.i., which were largely conserved between different time points (**Figure 6A-B, S5A-B**). These RNAs consisted mostly of host mRNAs, however we also noticed a robust and consistent enrichment for the RNA components of the mitochondrial ribosome (mito-ribosome, 12S and 16S RNAs) in both VeroE6 and Huh7.5 cells (**Figure 6A, 6B**).

**Figure 6:**
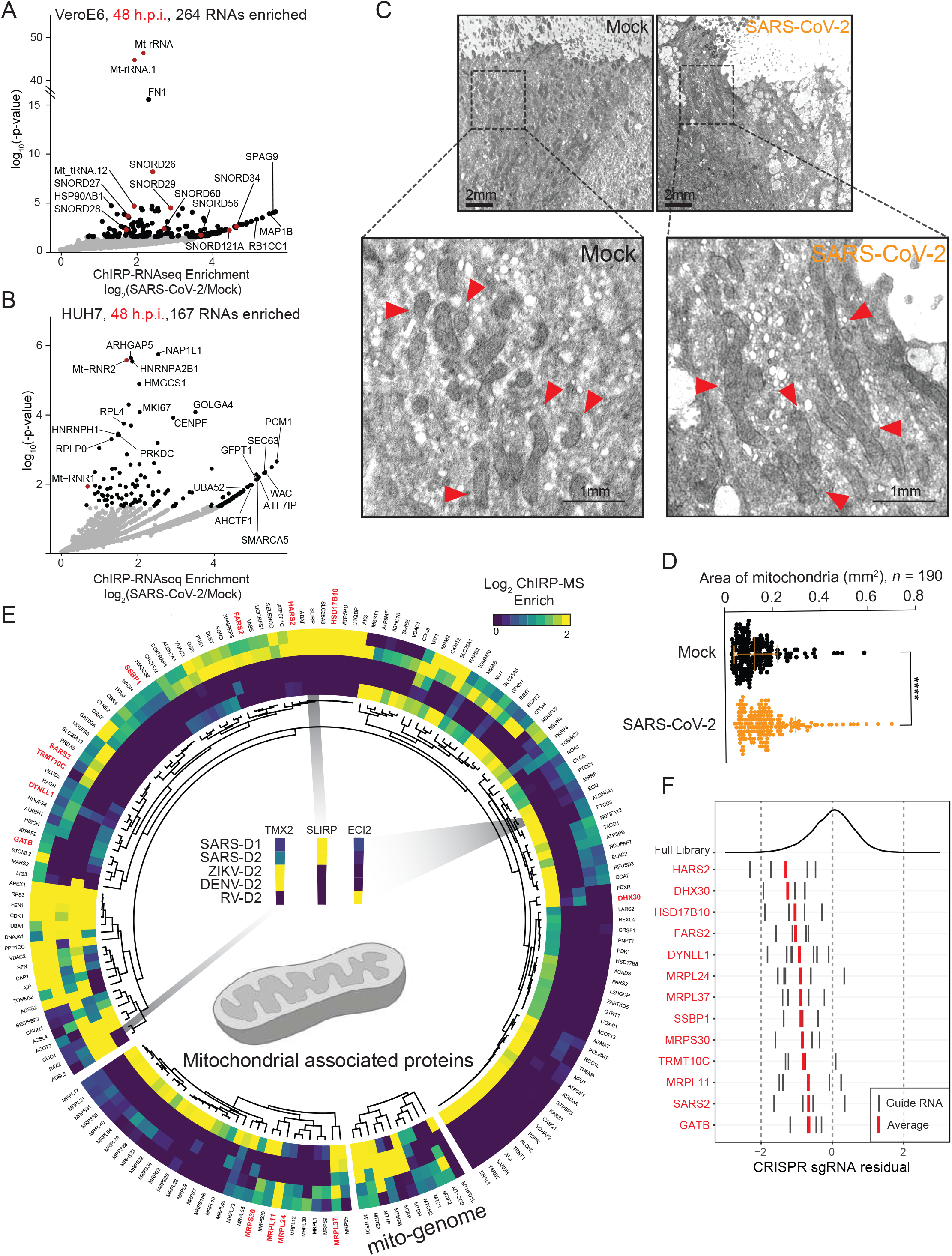
SARS-CoV-2 associated proteins and RNAs nominate the mitochondria in viral pathogenesis. **(A)** Enriched host RNAs after vRNA pulldown in VeroE6 cell line 48 h.p.i. **(B)** Enriched host RNAs after vRNA pulldown in Huh7.5 cell line 48 h.p.i. **(C)** Electron microscopy (EM) of HBEC cells uninfected (left, Mock) or infected by SARS-CoV-2 (right). Selected mitochondria indicated with arrowheads in the inset. **(D)** Quantification of mitochondria size by EM in five ciliated (infected) cells. **(E)** Mitochondrial proteins which are present in the expanded interactome of at least one virus and their conservation across viruses. Segments of the circle, from smallest to largest, correspond to proteins encoded by the mitochondrial genome, components of the mitochondrial ribosome, and proteins encoded by the nuclear genome which are localized or associated with the mitochondria. Enrichment (log_2_ FC) scale is capped at 2. Proteins which are significant hits in the CRISPR screen data in Figure 5 are indicated with red labels. **(F)** CRISPR sgRNA residuals for mitochondrially annotated CRISPR hits labeled in red in E.

Within the rest of the enriched RNAs, we found 8 C/D box snoRNAs (e.g. SNORD26, **Figure 6A**), which canonocially guide the deposition of 2’-O-methylation. We therefore asked if the proteins guided by C/D box snoRNAs were also enriched with vRNAs. There are at least eight RNA 2’-O-methyltransferases encoded in the human genome (Ayadi et al., 2019; Dimitrova et al., 2019), and we found that three of these are recovered in at least one of our ChIRP-MS datasets: FBL, MRM2, and MRM3 (**Figure S5C**). FBL is the canonically active in the nucleolus, while MRM2 and MRM3 have roles more biased for the mitochondria (Ayadi et al., 2019; Dimitrova et al., 2019). All three enzymes showed virus-specific patterns with MRM2 having among the highest enrichment values of any protein enriched in SARS-CoV-2 Huh7.5 D2 cells (**Figure 2B, S5C**). DENV and ZIKV were both more selective for FBL and MRM2, and RV selective for MRM3; however, the relative strength of these interactions were substantially weaker than what we observed for the interaction between MRM2 and SARS-CoV-2 (**Figure S5C**). Together, the association of snoRNAs and MRM2 with the SARS-CoV-2 vRNA suggests a specific interaction between SARS-CoV-2 and the host mitochondria with a possible role for 2’-O-methylation in SARS-CoV-2 infection.

The RNA- and protein-based associations with mitochondrial factors supports close physical contact between the mitochondria and the virus during infection. Indeed, Wu et al. recently reported that SARS-CoV-2 sgRNAs, particularly its 5’ untranslated region, contains sequence elements that strongly direct residency in mitochondria (Wu et al., 2020). Mitochondria are critical mediators of cellular homeostasis, we therefore hypothesized that viral infection may disrupt normal mitochondrial activities. Because the number and physical organization of mitochondria can directly indicate the function of these organelles, we set out to understand what, if any, changes occur during infection of human cells with SARS-CoV-2. We analyzed electron microscopy images of SARS-CoV-2 infected human bronchial epithelial cells (HBECs, 48 h.p.i.) with a focus on the mitochondria (**Figure 6C**, (Wei et al., 2020)). Upon visual inspection, the mitochondria within SARS-CoV-2-infected cells looked larger, which we confirmed by quantifying 190 mitochondria in five ciliated (infected) cells (**Figure 6C, 6D**). There was a significant increase in average mitochondrial size in cells infected with SARS-CoV-2 (**Figure 6D**), suggesting perturbed mitochondrial homeostasis.

To better understand the specificity and impact SARS-CoV-2 has on host mitochondria, we performed a focused reexamination of the ChIRP-MS data on mitochondrially-related proteins. We extracted ChIRP-MS enrichment values for human proteins associated with mitochondria (UniProtKB subcellular location of SL-0173), as well as mitochondrial-genome encoded proteins (**Figure 6E**). Overall, we found 162 (of 810 total, **Table S3**) mito-annotated proteins present in the expanded interactomes of at least one virus. Both DENV and ZIKV had relatively poor recovery of these proteins, with the exception of the mito-genome encoded proteins MT-AP, MT-MR6, MT-REX, MD-DH, MT-TP, and MT-HFD1 (**Figure 6E, bottom**). RV and SARS-CoV-2 both robustly bound to many of these factors but the particular associations are mostly non-overlapping (**Figure 6E**, examples are inset). This suggests that while vRNA may commonly associate with mitochondrial protein factors, there are virus-specific associations which may lead to differential pathological outcomes.

To assess the functional consequences of mitochondrial perturbation in the context of SARS-CoV-2 infection, we reanalyzed the CRISPR screen data focused on the same 810 mitochondrially-related proteins. We identified thirteen mitochondria-annotated proteins which were both present in the SARS-CoV-2 expanded interactome and significant hits in the genome-wide CRISPR screen (**Figure 6F**). Knockout of any of these thirteen genes individually caused increased cell death in the presence of infection (**Figure 6F**). Inspecting the genes, we observed an enrichment for those involved in protein synthesis by the mitochondrial ribosome including: components of the mitochondrial ribosome (MRPL11, MRPL24, MRPL37, and MRPS30), tRNA synthetases (FARS2, HARS2, and SARS2), an amidotransferase (GATB), a methyltransferase (TRMT10C), an RNA helicase (DHX30), and a DNA binding protein (SSBP1). Mapping these back to the ChIRP-MS enrichment, all of these host mitochondrial factors, except DHX30, exhibited specific or biased enrichment for SARS-CoV-2 RNA compared to that of other viruses (**Figure 6E, red labels**). Four components of the mitochondrial ribosome (MRPS2, MRPS5, MRPS25, and MRPS27, all distinct from the ChIRP-MS enriched mito-ribosomal proteins) have additionally been reported to interact with the viral protein nsp8, although the functional relevance was not determined or speculated upon at the time (Gordon et al., 2020). More recently, functional work (CRISPR screening in the Huh7.5 cell line) has independently validated the importance of all four of these proteins (Hoffmann et al., 2020). Altogether, these results point to a functionally relevant logic between viral RNAs and host mitochondria, and more generally suggest the mitochondria may be a major response organelle that protects from virus-induced death.

## Discussion

In summary, our results provide an RNA-centric view of the landscape of the host proteins and RNAs interacting with SARS-CoV-2 RNA during the course of infection. By integrating our analysis across time points, species, and other viruses, we identify shared and SARS-CoV-2-specific patterns of RNA-host protein interactions. In the context of the rapidly evolving literature on subcellular mechanisms of SARS-CoV-2 pathogenicity, the ChIRP-MS data provides an orthogonal but complementary resource to existing protein-protein interaction and phenotypic CRISPR screening studies (Gordon et al., 2020; Hoffmann et al., 2020; Wang et al., 2020; Wei et al., 2020). In particular, we find that the vRNA:host protein interface is largely distinct from that of viral and host proteins and nominates roles for previously unappreciated biological processes in SARS-CoV-2 infection. Our data also reveal unexpected aspects of virally encoded proteins. For example, many of the predicted viral RBPs were robustly captured by ChIRP in VeroE6 cells (e.g. nsp12 and nsp13); however, these were not enriched during infection in Huh7.5 cells. These cell type or species-specific associations could provide insights into differential virus susceptibility phenotypes, such as high viral titer and rapid cytotoxicity observed in VeroE6 cells, compared to Huh7.5 cells.

Comparisons of SARS-CoV-2 ChIRP-MS data to ChIRP-MS of three other positive-sense RNA viruses, and to genome-wide CRISPR screens, provided several new insights into the ‘molecular arms race’ that takes place between the virus and host. First, this analysis identified shared and unique strategies employed by viruses to hijack the host for trafficking and replication. For example, SARS-CoV-2 and flavivirus vRNAs both associate with the Rab GTPase proteins, RAB10 and RAB2A, which are involved in subcellular trafficking, and CRISPR perturbation revealed that these proteins are required for viral replication and virus-induced cell death (Gordon et al., 2020; Hoffmann et al., 2020). In contrast, despite the fact that both viral families depend on glycoproteins to produce infectious virions, there was a limited association of SARS-CoV-2 vRNA with the translational apparatus and the Sec/Translocon/OST complexes, compared to flaviviruses (Ooi et al., 2019). These data suggest that while both form membrane-enclosed replication complexes, flaviviruses may physically leverage the translocon complex, while SARS-CoV-2 leverages other domains of the ERGIC. Second, an unexpected finding from the intersection of ChIRP-MS and CRISPR datasets was that the vast majority (116/138) of vRNA-binding proteins were host-protective, rather than pro-viral factors. Most of these factors were commonly bound to multiple viral families, but we also found 31 SARS-CoV-2-specific factors. These results demonstrate that host cells deploy a broad and diverse array of proteins to physically recognize and counteract viral infection, and that these proteins are not limited to those with well-characterized viral recognition function, such as Toll-like receptors (TLRs) and RIG-I-like receptors (RLRs), but also extend to many other protein families with RNA-binding capacity. It is important to note that in this study, we relied on cell death-based CRISPR screens to assign functionality to vRNA-binding factors, but future screens focused on other aspects of the viral life cycle may identify additional functional aspects of these factors.

Finally, we identified a functional connection between SARS-CoV-2 vRNA and the mitochondria. Both RNA and protein components of the mitochondria were robustly captured with the SARS-CoV-2 vRNA in VeroE6 and Huh7.5 cells, suggesting a close physical interaction, and electron microscopy demonstrated changes in mitochondrial shape and size after infection. Interestingly, other viruses, including HIV have also been reported to physically enter the mitochondria, providing evidence that vRNA can gain access to the mitochondria during infection (Somasundaran et al., 1994). Mitochondria are central to the underlying health of a cell, play an active role in sensing and signaling during cellular stress and act as a hub for innate immune signaling. Indeed, we found that functional CRISPR perturbations of mitochondrial proteins revealed that many of these proteins were required for host survival. Based on our results, we propose that RNA viruses may follow a distinct logic when causing mitochondrial stress; that is, many viruses may interact with and perturb this organelle, but the precise manner in which stress is caused, and thus signaling occurs, is virus-specific. Along these lines, one of the top virus-specific hits in the Huh7.5 ChIRP-MS data was MRM2/FTSJ2, a mitochondrial localized 2’-O-methyltransferase. This is of particular interest due to the previous identification and characterization of FTSJ3/SPB1 as a factor that methylates the HIV RNA genome, which leads to pro-viral shielding of the HIV RNA from MDA5 recognition (Ringeard et al., 2019). Thus, while additional work is needed to define the mechanism of action of MRM2 in the context of SARS-CoV-2 infection, we hypothesize that it may play a pro-viral role. Altogether, this study provides an unbiased and comprehensive catalogue of functional SARS-CoV-2 RNA-host protein interactions, revealed a functional link between SARS-CoV-2 and the mitochondria, and may inform future studies to understand the mechanisms of viral pathogenesis and nominate strategies to combat the virus for therapeutic benefit.

## Supplementary Figure Legends

**Figure S1.**
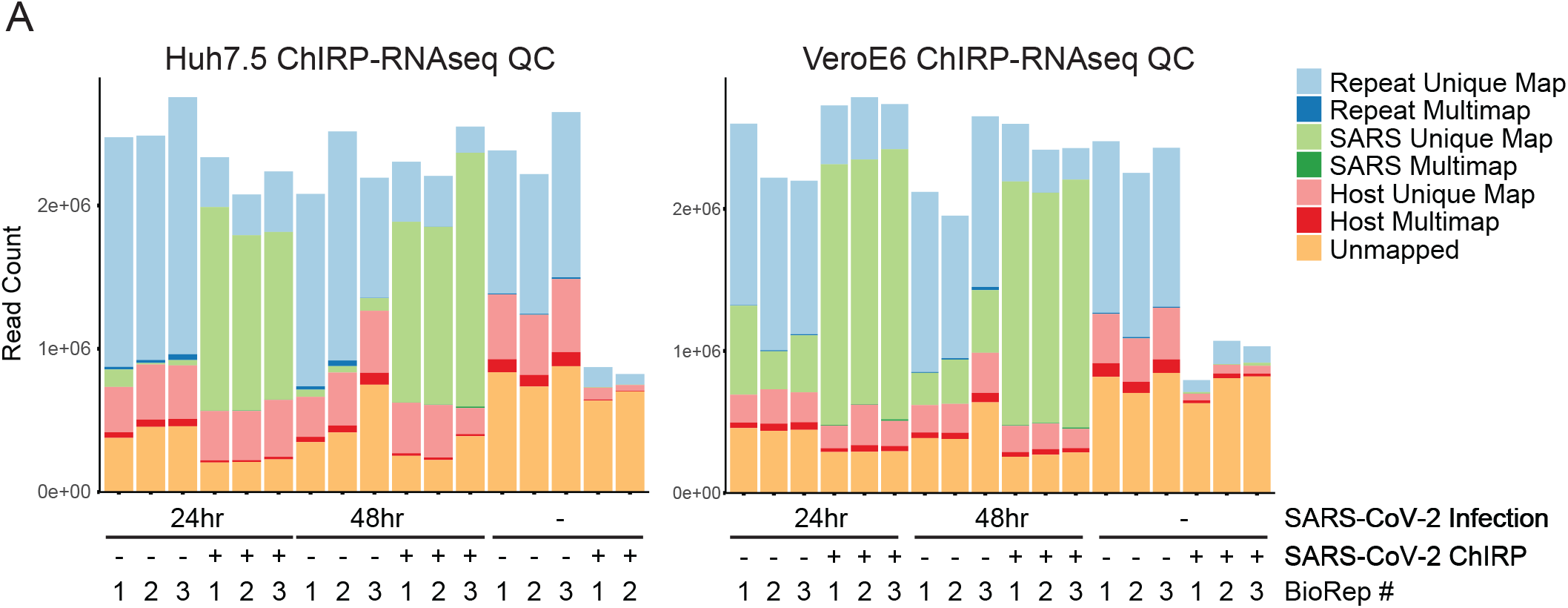
**related to Figure 1 (A)** Host and viral RNA-seq alignment statistics for all samples across Huh7.5 (left) and VeroE6 (right) cell lines.

**Figure S2.**
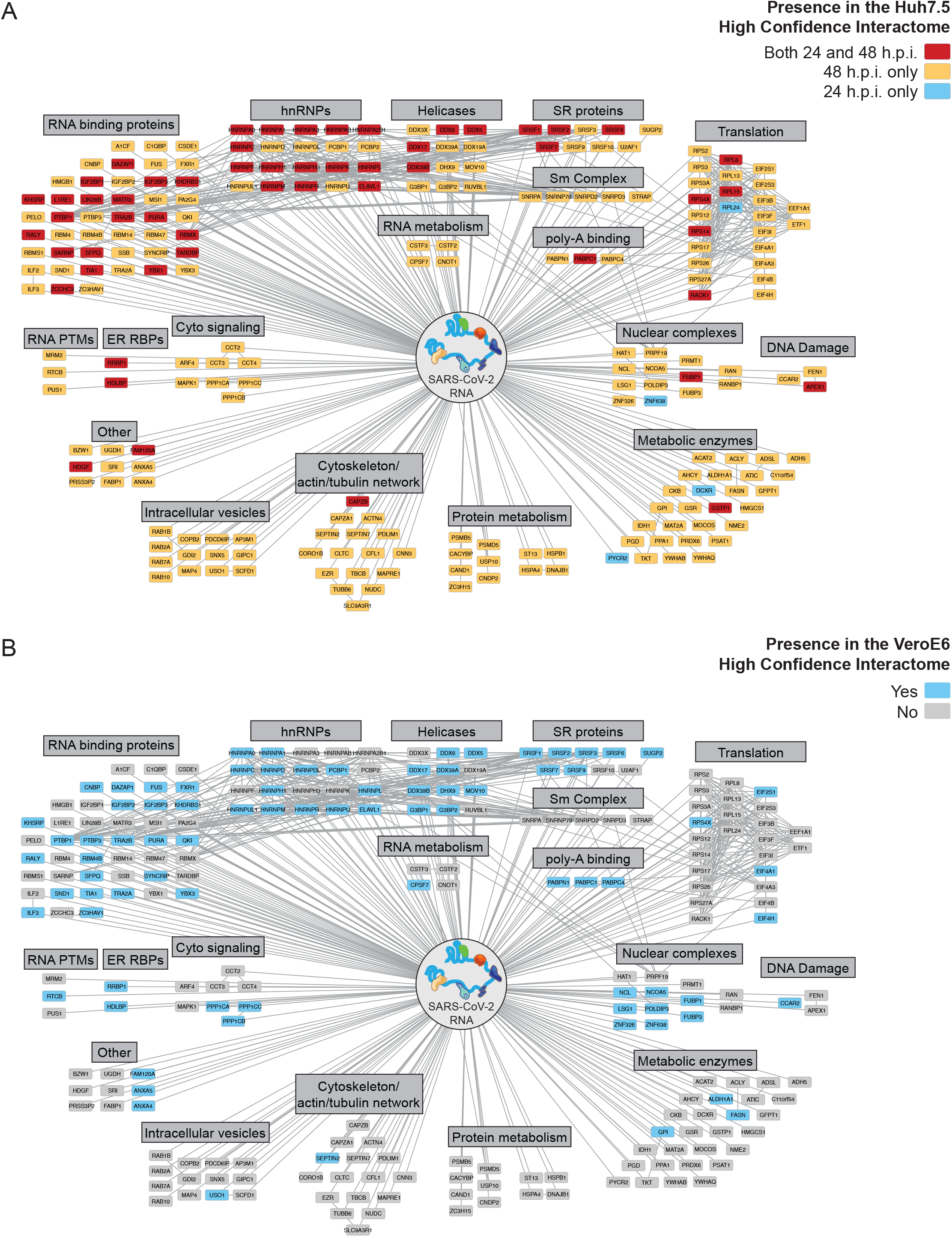
**related to Figure 2 (A)** High confidence SARS-CoV-2 human interactome network colored by time point (24 h.p.i., 48 h.p.i., or both). **(B)** High confidence SARS-CoV-2 human interactome network colored by species conservation.

**Figure S3.**
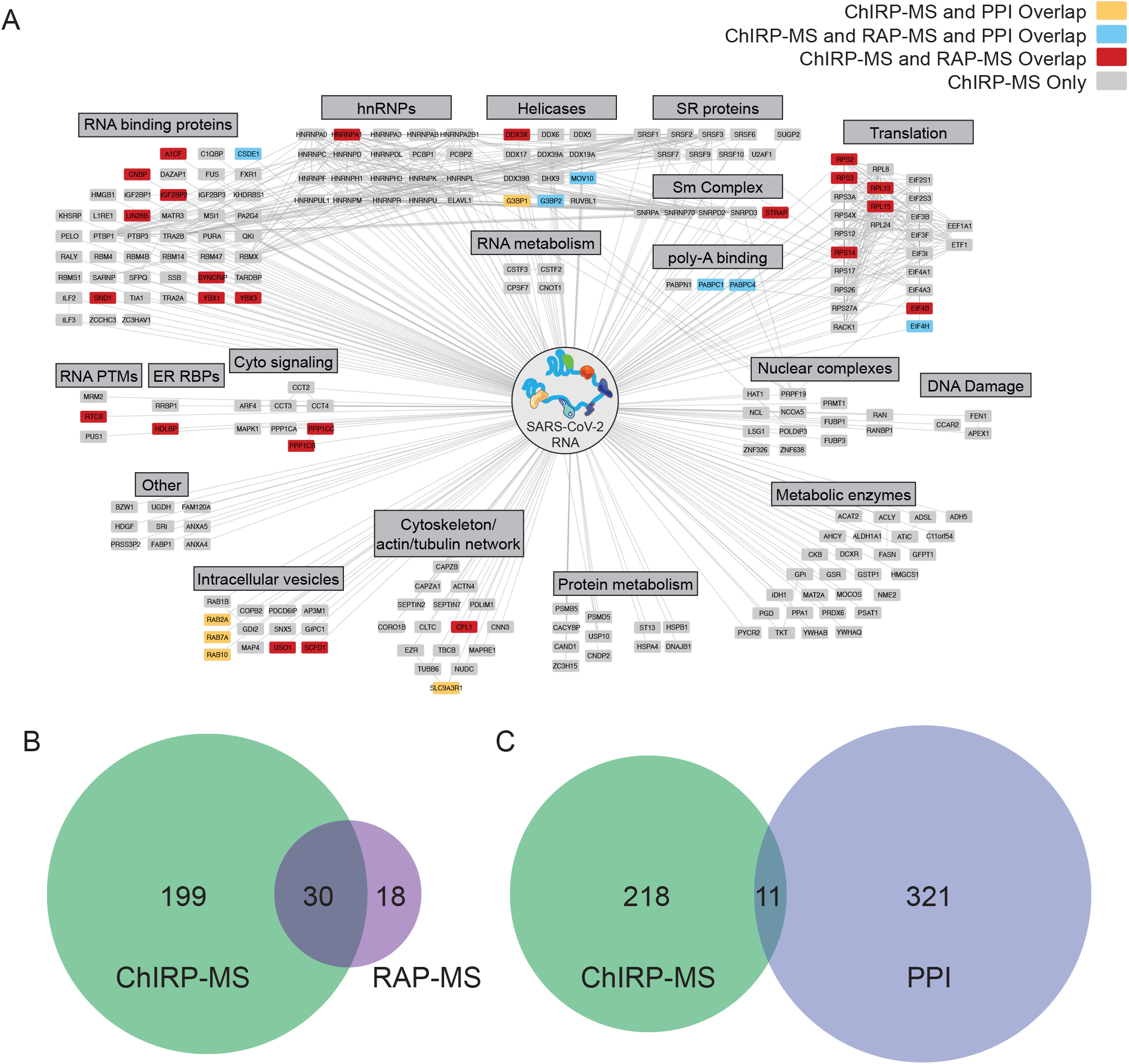
**related to Figure 2 (A)** Comparison of the high confidence SARS-CoV-2 RNA associated human proteome by RAP-MS (UV crosslinking) to that by formaldehyde crosslinking (ChIRP-MS) and comparison of the SARS-CoV-2 RNA associated proteome to the SARS-CoV-2 protein associated proteome (PPI). **(B)** Overlap of interactomes by RAP-MS and ChIRP-MS. **(C)** Overlap of interactomes by PPI and ChIRP-MS.

**Figure S4.**
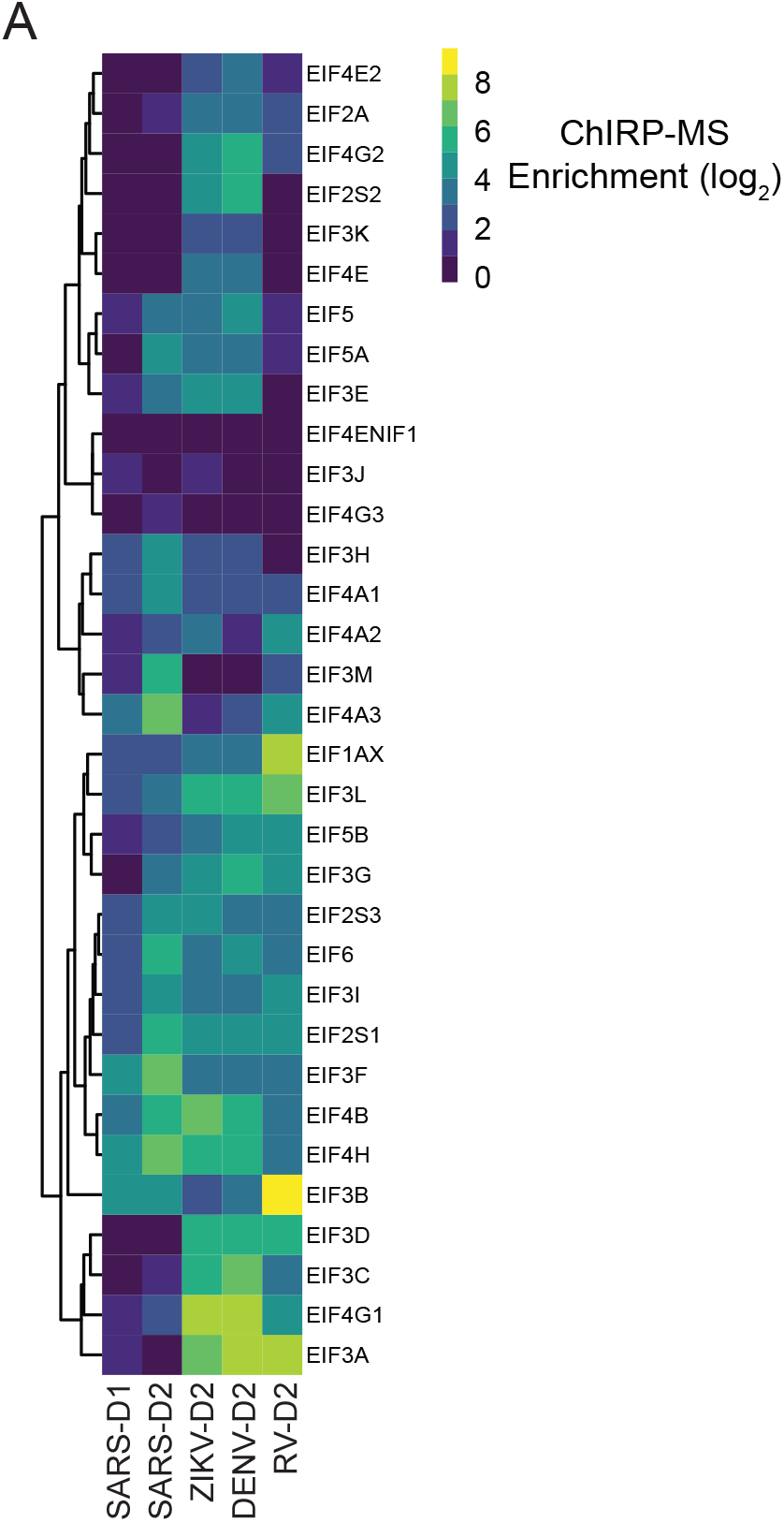
**related to Figure 4 (A)** Pan-viral comparison of associations with translation initiation (EIF) factors.

**Figure S5.**
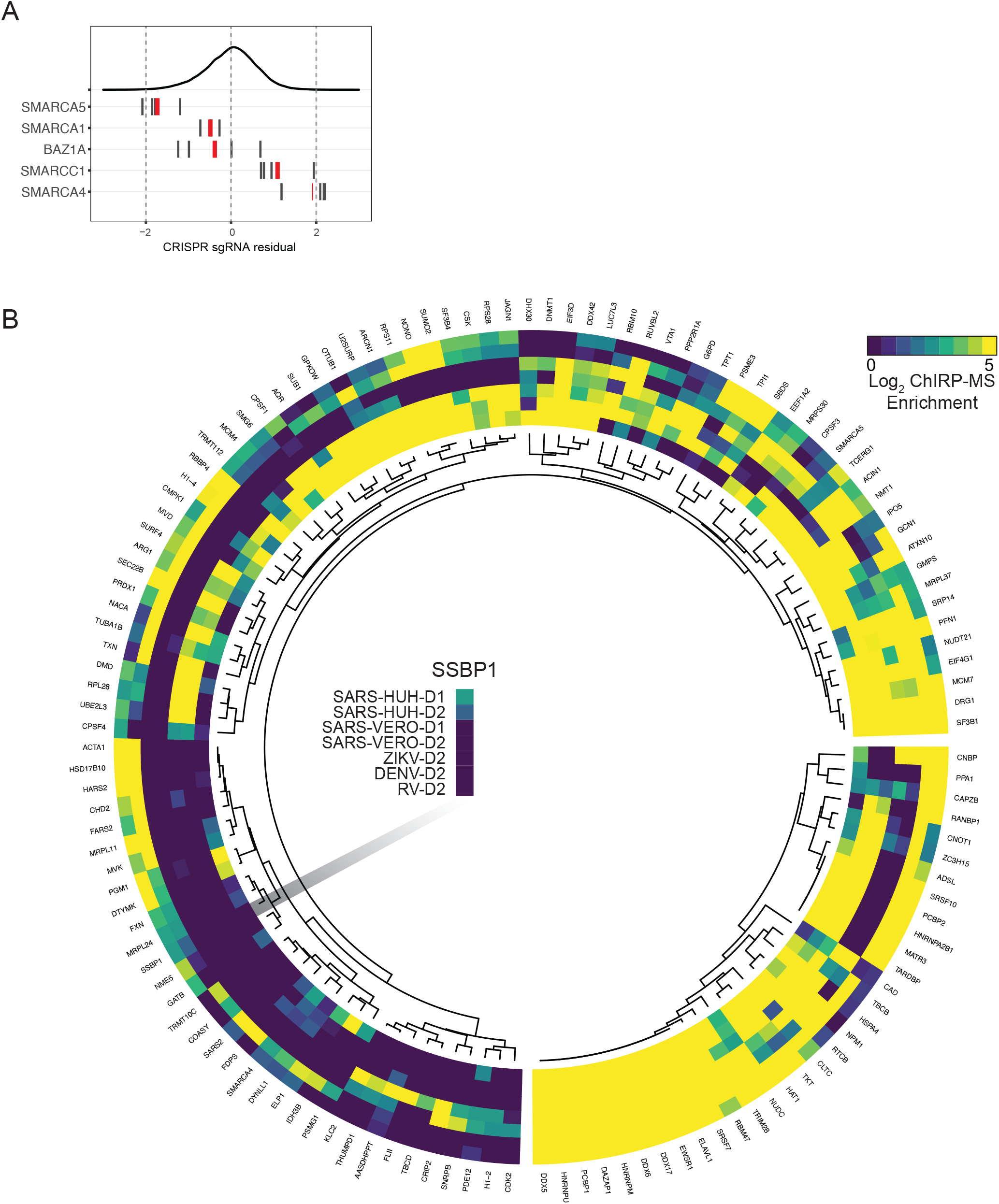
**related to Figure 5 (A)** CRISPR guide residuals for SWI/SNF related genes enriched in the ChIRP-MS dataset. **(B)** All genes which are significant hits in the CRISPR screen (fdr ≤ 0.05) and also present in the high confidence interactome (smaller segment) or the expanded interactome (larger segment) of SARS-CoV-2 and their association with other viruses. Scale is capped at 5.

**Figure S6.**
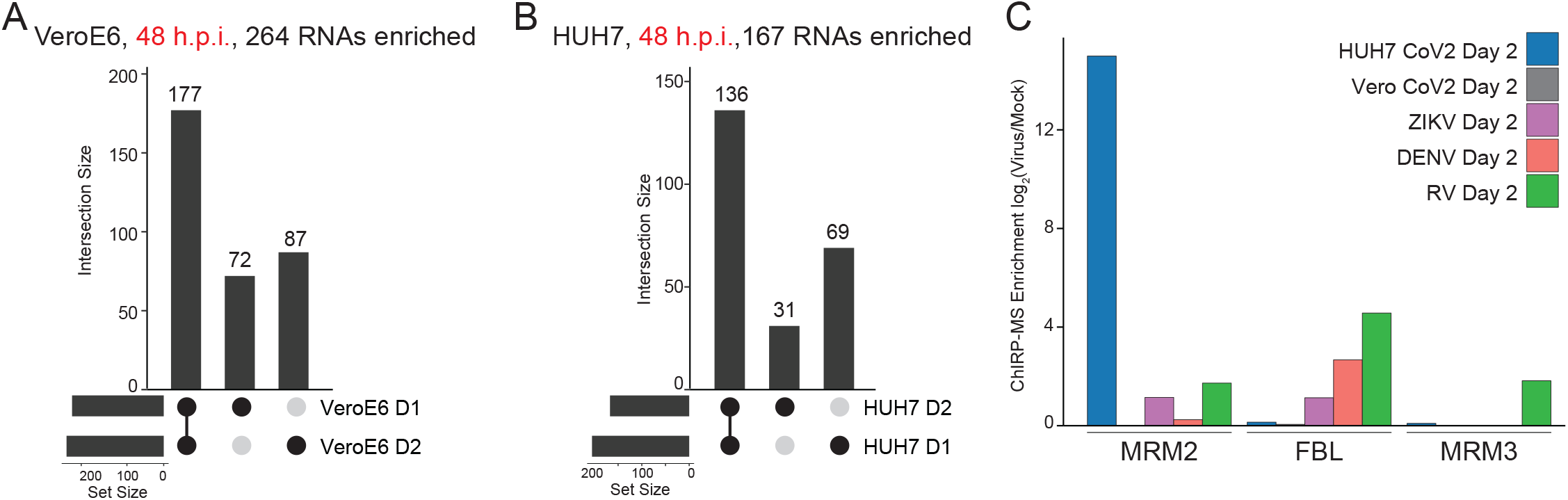
**related to Figure 6 (A)** Conservation between time points of SARS-associated host RNAs in the VeroE6 cell line. **(B)** Conservation between time points of SARS-associated host RNAs in Huh7.5 cell line. **(C)** ChIRP-MS enrichment of rRNA 2’-O-ribose methyltransferases across viruses.

## Table Descriptions

**Table S1:** Sequences of 108 biotinylated ChIRP probes.

**Table S2:** Mass spectrometry data for viral proteins and host proteins with mean enrichment ≥ 1 in SARS-CoV-2 ChIRP-MS datasets.

**Table S3:** Full data for all proteins in all datasets including pan-virus data, high confidence interactomes, expanded interactomes, CRISPR screen results, and presence in other datasets (such as UV-crosslinking data, PPI data).

## Methods

### Cell lines, SARS-CoV-2 infection, and cell processing

Vero-E6 and Huh7.5 cells were seeded at 1×10^6^ cells per T150 flask and were grown in Dulbecco’s Modified Eagle Medium (DMEM) supplemented with 10% heat-inactivated fetal bovine serum (FBS), and 1% Penicillin/Streptomycin. Three T150 flasks were assigned per condition: 0, 1, and 2 days post-infection (dpi). The next day, the media was removed, and cells were inoculated with SARS-CoV-2 isolate USA-WA1/2020 (BEI Resources #NR-48814) at MOI of 0.01. Flasks were incubated at 37°C for 1 hr with gentle rocking every 15 min. At 0, 1, and 2 dpi, supernatant from the flasks were discarded, and cells were washed with 1X PBS twice. 4 mL of 4% of paraformaldehyde was added on each of the flasks and incubated for 30 min at room temperature. Afterwards, cells were quenched with 250 µL of 2 M glycine (final concentration of 125 mM) for each flask. Cells were scraped, harvested in pre-weighed microcentrifuge tubes, and span at 1000 x g for 5 min at 4°C. All supernatants aspirated, and the final pellet were weighed. Cells were frozen at −80°C until used. All procedures with infectious virus were done at a Biosafety Level 3 (BSL3) laboratory and approved by the Yale University Biosafety Committee.

### Comprehensive identification of RNA binding proteins by mass spectrometry (ChIRP-MS)

SARS-CoV-2 targeting probes were designed online (https://www.biosearchtech.com/stellaris), with repeat masking setting of 3 and even coverage of the whole transcript. Full probe sequences available in **Table S1**. Oligos were synthesized with a 3′ biotin-TEG modification at Stanford Protein and Nucleic Acid Facility.

ChIRP-MS was performed largely as described in (Chu et al., 2015). Cells were cultured, infected, and crosslinked as described above in the BSL3 facility. Lysate was generated by resuspending cell pellets in 1 mL lysis buffer (50 mM Tris-HCl pH 7.0, 10 mM EDTA, 1% SDS) per 100 mg of cell pellet weight (∼100µL pellet volume). Lysates were sonicated using a focused-ultrasonicator (Covaris, E220) until the average RNA length was ∼500 nucleotides as determined by agarose gel analysis and stored at −80°C. Stored lysates were thawed on ice and prepared for pre-clearing. Precleared was achieved by adding 30 μL washed MyOne C1 beads per mL of lysate at 37°C for 30 minutes on rotation. Preclearing beads were removed twice from lysate using a magnetic stand; for this and all subsequent magnetic stand steps allow for > 1 minutes of separation before removing any supernatant. Next, for every 1 mL of sonicated lysate 2 mL of ChIRP hybridization buffer (750 mM NaCl, 1% SDS, 50 mM Tris-HCl pH 7.0, 1 mM EDTA, 15% formamide; made fresh) and 2.5 µL of 100 µM ChIRP Probe Pools were added per mL of lysate. ChIRP Probe Pools (**Table S1**) were composed of an equimolar mix of 108 antisense oligos. For each biological triplicate, a total of 7 mL of sonicated cell lysate was used. Hybridization took place on rotation for 16 hours at 37°C. Subsequently, 250 µL of washed MyOne C1 beads per mL of lysate were added to each sample and incubated on rotation for 45 minutes at 37°C. Enriched material was collected on the beads with a magnetic stand, and beads were washed 5x 2 minutes in 1 mL of ChIRP Wash Buffer (2x NaCl-Sodium Citrate (SSC, ThermoFisher Scientific), 0.5% SDS) at 37°C. After washing, 1% of each sample was saved for RNA extraction and RNA-seq library preparation (below). To elute enriched proteins, beads were collected on magnetic stand, resuspended in ChIRP biotin elution buffer (12.5 mM biotin, 7.5 mM HEPES, pH 7.9, 75 mM NaCl, 1.5 mM EDTA, 0.15% SDS, 0.075% sarkosyl, and 0.02% Na-Deoxycholate), mixed at 25°C for 20 minutes on rotation and at 65°C for 15 minutes shaking. Eluent was transferred to a fresh tube, and beads were eluted again. The two eluents were pooled (∼1200 µL), and residual beads were removed again using the magnetic stand. 25% total volume (300 µL) trichloroacetic acid was added to the clean eluent, vortexed, and then samples were placed at 4°C overnight for precipitation. The next day, proteins were pelleted at 21,000 rcf at 4°C for 60 minutes. Supernatant was carefully removed and protein pellets were washed once with ice-cold acetone. Samples were spun at 21,000 rcf at 4°C for 5 minutes. Acetone supernatant was removed, tubes briefly centrifuged again and, after removal of residual acetone, were left to air-dry on the bench-top. Proteins were then solubilized in 1x LDS Buffer in NT2 with 20 mM DTT and boiled at 95°C for 30 minutes with occasional mixing for reverse-crosslinking.

Protein samples were size-separated on bis-tris SDS-PAGE gels (Bio-Rad), and the gel was fixed and stained with the Colloidal Blue Staining Kit (ThermoFisher Scientific) as per the manufacturer’s instructions. Each ChIRP-MS experiment (1 lane in the gel) was cut into 2 slices from the SDS-PAGE and prepared independently. Gel slices were prepared for mass spectrometry by rinsing sequentially in 200 µL HPLC-grade water, 100% Acetonitrile (ACN, ThermoFisher Scientific), 50 mM Ammonium Bicarbonate (AmBic). Samples were reduced by adding 200 µL of 5 mM DTT in 50 mM AmBic and incubating at 65°C for 35 minutes. The reduction buffer was discarded, and samples were cooled to room temperature. Alkylation was achieved by adding 200 µL of 25 mM iodoacetamide in 50 mM AmBic for 20 minutes at 25°C in the dark. The alkylation buffer was discarded, samples were rinsed once in 200 µL 50 mM AmBic, and then they were washed twice for 10 minutes each in 200 µL of freshly prepared 50% ACN in 50 mM AmBic. After each wash, the supernatant was discarded, and after all washes, samples were dried for 3 hours using a SpeedVac. Once dry, proteins were digested by adding 100 ng of trypsin in 200 µL of 50 mM AmBic for 16 hours at 37°C. Samples were subsequently acidified by adding formic acid to a final concentration of 1% and incubating at 37°C for 45 minutes. Finally, samples were desalted using HyperSep Filter Plates with a 5-7 µL bed volume (ThermoFisher Scientific) following the manufacturer’s instructions. Samples were eluted twice in 100 µL 80% ACN in 0.2% formic acid, dried on a SpeedVac, and resuspended in 10 µL 0.1% formic acid for mass spectrometry analysis.

All samples were resuspended in 10 μL 0.2% formic acid in water and 4 μL were injected on column for each sample. Peptides were separated over a 50 cm EasySpray reversed phase LC column (75 µm inner diameter packed with 2 μm, 100 Å, PepMap C18 particles, Thermo Fisher Scientific). The mobile phases (A: water with 0.2% formic acid and B: acetonitrile with 0.2% formic acid) were driven and controlled by a Dionex Ultimate 3000 RPLC nano system (Thermo Fisher Scientific). An integrated loading pump was used to load peptides onto a trap column (Acclaim PepMap 100 C18, 5 um particles, 20 mm length, ThermoFisher) at 5 µL/min, which was put in line with the analytical column 6 minutes into the gradient for the total protein samples. Gradient elution was performed at 300 nL/min. The gradient increased from 0% to 5% B over the first 6 minutes of the analysis, followed by an increase from 5% to 25% B from 6 to 86 minutes, an increase from 25% to 90% B from 86 to 94 minutes, isocratic flow at 90% B from 94 to 102 minutes, and a re-equilibration at 0% for 18 minutes for a total analysis time of 120 minutes. Precursors were ionized using an EASY-Spray ionization source (Thermo Fisher Scientific) source held at +2.2 kV compared to ground, and the column was held at 45 °C. The inlet capillary temperature was held at 275 °C, and the RF lens was held at 60%. Survey scans of peptide precursors were collected in the Orbitrap from 350-1350 Th with an AGC target of 1,000,000, a maximum injection time of 50 ms, and a resolution of 120,000 at 200 m/z. Monoisotopic precursor selection was enabled for peptide isotopic distributions, precursors of z = 2-5 were selected for data-dependent MS/MS scans for 2 seconds of cycle time, and dynamic exclusion was set to 45 seconds with a ±10 ppm window set around the precursor monoisotope. An isolation window of 1 Th was used to select precursor ions with the quadrupole. MS/MS scans were collected using HCD at 30 normalized collision energy (nce) with an AGC target of 50,000 and a maximum injection time of 54 ms. Mass analysis was performed in the Orbitrap with a resolution of 30,000 at 200 m/z and an automatically determined mass range.

FASTA sequences of the human proteome (Uniprot: UP000005640) were used and FASTA sequences of the viral proteins from SARS-CoV-2 (Uniprot: P0DTC1, P0DTD1, P0DTC2, P0DTC3, P0DTC4, P0DTC5, P0DTC6, P0DTC7, P0DTD8, P0DTC8, P0DTC9, P0DTD2, P0DTD3, A0A663DJA2), DENV (Uniprot: A0A173DS53), ZIKV (Uniprot: A0A140D2T1), RV (Uniprot: P03303) were appended to the end of the human proteome reference file. For the VeroE6 reference: GreenMonkey (Chlorocebus sabaeus, Uniprot: UP000029965). This concatenated file was used to search the ChIRP-MS data with MaxQuant with the following parameters: semi-specific cleavage specificity at the C-terminal site of R and K allowing for 2 missed cleavages. Mass tolerance was set at 12 ppm for MS1s, 0.4 for MS2s. Methionine oxidation, asparagine deamidation, and N-term acetylation were set as variable modifications. Cysteine carbamidomethylation was set as a fixed modification. Label-free quantitation values from MaxQuant were imported into R for downstream analysis. To define significantly enriched SARS-associated protein sets, R package ‘Differential Enrichment analysis of Proteomics data’ (DEP) was used. Filtering, normalization, and imputation were performed on MaxQuant outputs using the DEP default workflow. Enriched protein sets were defined using cutoffs log_2_ fold change > 0 and adjusted p-value <= 0.05, comparing infected cells after SARS RNA pulldown to identically treated uninfected (mock) cells.

### ChIRP-RNA-seq and analysis

Input lysate samples and enriched RNA samples (1% of the ChIRP sample) were first digested of their cellular proteins which also acts to effectively reverse the formaldehyde crosslinking. RNA samples were brought to 50 μL with 1x PBS and 5uL Proteinase K (Thermo Fisher Scientific) and incubated at 55C for 30 minutes. RNA was cleaned using the Zymo Clean and Concentrate 5 column (Zymo Research) and eluted in 2x 20 μL (final 40 μL). DNA was removed by adding 2 μL DNaseI and 5 μL 10x DNase buffer (NEB) to the purified RNA and incubated at 37C for 30 minutes. The RNA was cleaned up as above with the Zymo Clean and Concentrate 5 column but eluted 2x 10 μL (final 20 μL). To construct RNA seq libraries, TAKARA Bio SMART-Seq Stranded Kit User Manual (TAKARA Bio) was used with the following modifications. Up to 5 ng RNA was reverse-transcribed and amplified by PCR following the SMART-seq protocol. To increase cDNA yield and detection efficiency, we started from first-strand cDNA synthesis without fragmentation. The number of PCR1 cycles was 5. We purified the cDNA product with 50 μL AMPure beads (1:1 ratio) and eluted into 20 μL water. Then the 20 μL purified cDNA was used as input for the final RNA-Seq library amplification. To reduce the amount of primer dimer artifacts, we purified the RNA-Seq library with 90 μL AMPure beads (x0.9 selection) and eluted into 20 μL water. Sequencing was performed using the Nextseq 500/550 Sequencing system (Illumina) with 2 x 75 bp paired-end reads and 2 x 8 bp index reads.

Adapters were automatically detected and trimmed using fastp^5^. Host genomes (for homo sapiens and chlorocebus sabaeus) were obtained from Ensembl along with annotation (gtf) files for use with feature counts. The SARS-CoV-2 genome was obtained from NCBI. Hisat2 was used to index all genomes and align reads^6^. Fastq files were initially aligned to a file of known “repeat” sequences--specific sequences which are present in multiple locations in the genome and which can cause a high percentage of multi-mapped reads. Remaining reads were then aligned to the SARS-CoV-2 genome. SARS-CoV-2 genome coverage was visualized in the Integrative Genomics Viewer to assess pulldown efficiency. Remaining reads were then aligned to the host genome and reads overlapping genomic features (genes) were quantified using the featureCounts command line utility^7^. Aggregated counts matrices were loaded into DESeq2 for normalization and differential gene expression analysis^8^. For GO term analysis, expanded interactomes of each virus were annotated with the DAVID Bioinformatics Resource (Huang et al., 2009a, 2009b). Annotations for Cellular Components, Binding Proteins, and Protein Domains were used to compute enrichments for each expanded interactome.

### Electron Microscopy

Samples were from Wei et al. 2020 and prepared in the following way: HBECs were fixed using 2.5% glutaraldehyde in 0.1M phosphate buffer, osmicated in 1% osmium tetroxide, and dehydrated in ethanol. During dehydration, 1% uranyl acetate was added to the 70% ethanol to enhance ultrastructural membrane contrast. After dehydration the cells were embedded in Durcupan and ultrathin sections were cut on a Leica Ultra-Microtome, collected on Formvar-coated single-slot grids, and analyzed with a Tecnai 12 Biotwin electron microscope (FEI). ImageJ software was used to measure mitochondrial area.

## Acknowledgement

We thank Nicholas Riley for designing the Mass Spec methods and operating the instrument, Chris Richards and Chris Lapointe for helpful discussions, John Doench, Ruth Hanna, and Peter DeWeirdt for assistance analyzing CRISPR data, and Sigrid Knemeyer and Christine Shan at SciStories LLC for illustrations. This work was supported by the National Institutes of Health grants K08CA230188 (A.T.S.), K08AI128043 (C.B.W.), R01AI140186S (J.E.C), and R01AI141970 (J.E.C.), the Burroughs Wellcome Fund Career Award for Medical Scientists (R.A.F., A.T.S., C.B.W.), the Bill and Melinda Gates Foundation (A.T.S.), the Ludwig Family Foundation (C.B.W.), a Stanford ChEM-H COVID-19 award (A.T.S.), and Fast Grants from Emergent Ventures (A.T.S., C.B.W.). R.A.F. was supported by Damon Runyon Cancer Research Foundation DRG-2286-17. J.A.B and K.R.P. were supported by Stanford Graduate Fellowships. J.A.B. was supported by the National Science Foundation Graduate Research Fellowship under Grant No. DGE-1656518. A.T.S. was supported by a Parker Bridge Scholar Award from the Parker Institute for Cancer Immunotherapy and a Cancer Research Institute Technology Impact Award. J.E.C. was supported by the Burroughs Wellcome Investigators in the Pathogenesis of Infectious Disease Award. H.Y.C was supported by the Pershing Square Foundation and RM1-HG007735. C.R.B. and H.Y.C. are Investigators of the Howard Hughes Medical Institute. The sequencing data was generated at the Stanford Functional Genomics Facility with instrumentation purchased with NIH grants S10OD018220 and 1S10OD021763.

## Author contributions

R.A.F., M.R.M., C.B.W., and A.T.S. conceived the study. R.A.F., Y.Q., C.O.S. performed SARS-CoV-2 and ChIRP experiments. R.A.F., J.A.B., A.L., K.R.P. performed analysis of ChIRP data. J.W., M.M.A., and C.B.W. performed and analyzed CRISPR screens. Y.Y. and T.H. performed mitochondrial imaging experiments. R.A.F., H.Y.C., T.H., J.E.C., C.R.B., C.B.W., and A.T.S. oversaw and guided experiments and analysis. R.A.F., J.A.B., C.B.W., and A.T.S. drafted the manuscript and all authors reviewed and provided comments on the manuscript.

